# Cytoplasmic mRNA granules regulate cell fate decisions during PINK1/Parkin mitophagy

**DOI:** 10.64898/2026.02.13.705862

**Authors:** Taiki Baba, Akimi Inoue, Yuya Nagahata, Hibiki Tsutsumi, Jun Takouda, Rena Onoguchi-Mizutani, Nobuyoshi Akimitsu, Susumu Tanimura, Kohsuke Takeda

## Abstract

Mitophagy is generally considered to promote cell survival by removing damaged mitochondria in response to mitochondrial stress, whereas apoptosis occurs during prolonged stress. However, the mechanisms that determine cell survival and cell death under these stress conditions remain poorly understood. Here, we showed that cytoplasmic mRNA granules, designated as mitophagy-induced mRNA granules (mitoRGs), were formed transiently and played an important role in cell fate decisions during PINK1/Parkin-dependent mitophagy. Although some components, such as G3BP1, were shared with stress granules (SGs), mitoRGs were distinct from SGs because mitoRG assembly required the mitochondrial protein phosphatase PGAM5. In response to mitochondrial stress, PGAM5 was released into the cytosol from mitochondria and incorporated into mitoRGs, but was then released back into the cytosol during mitoRG disassembly following prolonged mitochondrial stress, corresponding with the induction of apoptosis. Impairment of mitoRG assembly through G3BP1 depletion sensitized cells to apoptosis during mitophagy in a PGAM5-dependent manner. These results suggest that mitoRGs regulate cell fate decisions by spatiotemporally controlling PGAM5 and its pro-apoptotic activity during PINK1/Parkin mitophagy.

## Introduction

Mitochondria are persistently exposed to metabolic fluctuations, reactive oxygen species, and environmental insults, necessitating robust quality-control mechanisms to maintain cellular homeostasis. Failure of mitochondrial surveillance compromises bioenergetic function and stress signaling, thereby contributing to a broad spectrum of human diseases, including neurodegenerative disorders and cancer (Liu *et al*, 2024). To counteract mitochondrial dysfunction, cells have evolved specialized signaling pathways that sense mitochondrial damage and initiate adaptive responses (Eckl *et al*, 2021). Among these, the PINK1/Parkin pathway represents the central axis of mitochondrial quality control, mediating the selective autophagic elimination of damaged mitochondria through mitophagy (Narendra & Youle, 2024). Following the loss of mitochondrial membrane potential, the mitochondrial serine/threonine kinase PINK1 accumulates on the outer mitochondrial membrane, activates the E3 ubiquitin ligase Parkin, and triggers extensive ubiquitination of mitochondrial surface proteins, thereby recruiting the autophagy machinery (Narendra *et al*, 2010, 2008; Matsuda *et al*, 2010).

PINK1/Parkin-dependent mitophagy (PINK1/Parkin mitophagy) is generally considered to be a cytoprotective process that alleviates mitochondrial stress and promotes cell survival (Wu *et al*, 2015; Exner *et al*, 2007; Chen & Dorn, 2013); however, increasing evidence indicates that this pathway can be repurposed to promote apoptotic cell death in the event of sustained or excessive mitochondrial damage. In these contexts, Parkin-driven remodeling of the mitochondrial outer membrane and proteasomal degradation of pro-survival factors lead to cytochrome c release and caspase activation (Quarato *et al*, 2023; Carroll *et al*, 2014). These observations suggest that the PINK1/Parkin pathway acts as a critical decision node that interprets mitochondrial stress signals and determines if cells should engage adaptive survival programs or commit to apoptosis. Despite this conceptual framework, the molecular mechanisms governing the binary cell fate decisions downstream of mitochondrial damage remain unclear.

Post-transcriptional gene regulation has emerged as an important, but underappreciated, layer of cellular stress adaptation, shaping the integration of stress signals to control survival and death outcomes (Pizzinga *et al*, 2020). The dynamic assembly of cytoplasmic ribonucleoprotein condensates, most notably stress granules (SGs), is a hallmark of this regulatory layer, transiently sequestering translationally repressed mRNAs and RNA-binding proteins under diverse stress conditions (Marcelo *et al*, 2021). SGs are thought to promote cell survival via these mechanisms under adverse conditions (Paget *et al*, 2023; Arimoto *et al*, 2008; Fujikawa *et al*, 2023). Although SG biology has been studied extensively in the context of oxidative and proteotoxic stress, whether or not mitochondrial stress, particularly that associated with PINK1/Parkin mitophagy, engages analogous RNA-based regulatory mechanisms remains largely unexplored.

The mitochondrial serine/threonine phosphatase PGAM5 has emerged as a multifunctional regulator of mitochondrial stress responses, with reported roles in mitophagy, mitochondrial dynamics, and programmed cell death (Qi *et al*, 2025; Takeda *et al*, 2009). PGAM5 is anchored to the inner mitochondrial membrane via its transmembrane domain and undergoes regulated intramembrane proteolysis by presenilin-associated rhomboid-like protein (PARL) and overlapping activity with m-AAA protease (OMA1) in response to mitochondrial depolarization (Wai *et al*, 2016; Sekine *et al*, 2012). The cleaved form of PGAM5, including the phosphatase domain, is released from mitochondria into the cytosol under conditions that trigger PINK1/Parkin mitophagy (Yamaguchi *et al*, 2019; Bernkopf *et al*, 2018). PARL-mediated PGAM5 processing is linked to apoptotic signaling following carbonyl cyanide m-chlorophenylhydrazone (CCCP)-induced mitochondrial depolarization, suggesting that cytosolic PGAM5 functions as a pro-apoptotic effector during mitochondrial stress (Saita *et al*, 2017; Zhuang *et al*, 2013). However, the mechanisms by which PGAM5 activity is spatially and temporally regulated during PINK1/Parkin mitophagy to balance survival and death signaling remain unclear. Notably, we previously showed that mitochondria-derived PGAM5 translocated into the nucleus during PINK1/Parkin mitophagy, where it dephosphorylated nuclear factors involved in mRNA metabolism, thereby directly coupling mitochondrial stress to post-transcriptional gene regulation (Baba *et al*, 2021). These findings suggest that PGAM5 acts as a stress-responsive signaling molecule that coordinates mitochondrial dysfunction with RNA regulatory programs across different subcellular compartments.

In the current study, we identified a previously unrecognized class of cytoplasmic mRNA granules that assembled transiently during PINK1/Parkin mitophagy, which we termed mitophagy-induced RNA granules (mitoRGs). We demonstrated that mitoRGs were similar to SGs in that both required the RNA-binding protein G3BP1 as an essential nucleator, but were also distinct from SGs because, unlike SGs, their assembly required PGAM5 and was independent of eukaryotic translation initiation factor 2α (eIF2α)-mediated translational arrest. PGAM5 released from mitochondria was incorporated into mitoRGs and finally released back into the cytosol during mitoRG disassembly, suggesting that mitoRGs act as a buffering platform that transiently sequesters PGAM5 in the cytoplasm. Together with the finding that impaired mitoRG formation led to diffuse cytosolic distribution of PGAM5 and early induction of apoptosis depending on PGAM5 activity, mitoRGs appear to serve as an RNA-based regulatory layer linking mitochondrial quality control to cell fate decisions by spatiotemporally controlling PGAM5 during PINK1/Parkin mitophagy.

## Results

### Cytoplasmic mRNA granules are transiently formed depending on PGAM5 during PINK1/Parkin mitophagy

We previously found that PGAM5 dephosphorylated nuclear proteins involved in mRNA metabolism during PINK1/Parkin mitophagy (Baba *et al*, 2021). To investigate the significance of this molecular behavior, we monitored intracellular mRNA dynamics during mitophagy using poly(A)^+^ RNA fluorescence *in situ* hybridization (FISH) in HeLa cells stably expressing hemagglutinin (HA)-tagged Parkin (Parkin-HeLa cells). Mitochondrial uncouplers, such as CCCP, can effectively induce mitophagy in these cells. The proportion of cytosolic to nuclear mRNA increased after mitophagy induction by CCCP treatment (**Fig. 1A, B**), suggesting that mRNA export from the nucleus to the cytosol is promoted at a relatively early stage of mitophagy. Notably, the increase in cytosolic mRNA was accompanied by the formation of granule-like puncta in the cytoplasm, which gradually enlarged from 2–8 h after CCCP treatment (**Fig. 1A–D**). Three-dimensional reconstruction indicated that these mitophagy-induced mRNA puncta were spherical granules (**Fig. S1A**). Given that CCCP functions as a protonophore and affects a range of biological processes extending beyond those occurring within mitochondria, we treated cells with oligomycin A and antimycin A (OA), which block mitochondrial ATP synthase and the mitochondrial respiratory chain complex III, respectively, and have been shown previously to induce autophagy (Lazarou *et al*, 2015). Treatment with OA also induced mRNA granules in the cytoplasm (**Fig. S1B**). These results suggest that mitochondrial stress is responsible for the formation of cytoplasmic mRNA granules after induction of PINK1/Parkin mitophagy. To the best of our knowledge, this process has not been reported previously, and we therefore named these granules mitophagy-induced cytoplasmic mRNA granules (mitoRGs).

**Figure 1.**
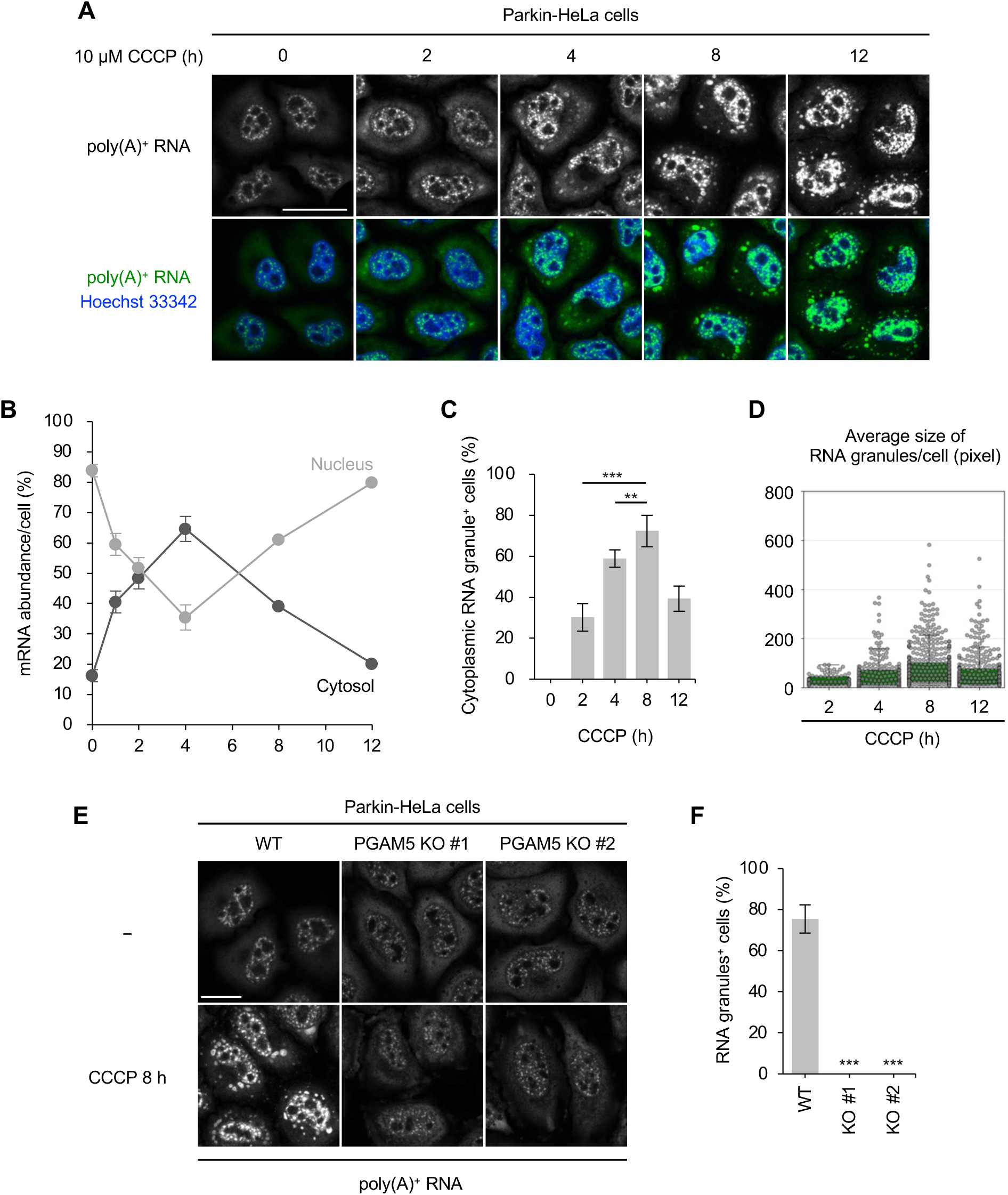
Cytoplasmic mRNA granules are transiently formed depending on PGAM5 during PINK1/Parkin mitophagy. (**A**) Poly(A)^+^ RNA-FISH of Parkin-HeLa cells treated with 10 μM CCCP for 2, 4, 8, and 12 h and counterstained with Hoechst 33342. Scale bar = 20 μm. (**B**) Quantification of proportion of mRNA in the cytoplasm and nucleus in (A). (**C**) Quantification of proportion of cytoplasmic RNA granule^+^ cells in (A). Tukey’s multiple-comparison test, **P < 0.01, ***P < 0.001. Error bars indicate mean ± SE. N = 3 independent experiments. (**D**) Average size of cytoplasmic RNA granules per cell (pixels) in (A). Two hundred cells per group were analyzed. (**E**) Parkin-HeLa cells (WT) and two lines of PGAM5 KO Parkin-HeLa cells (PGAM5 KO #1 and #2) were treated with 10 μM CCCP for 8 h and subjected to poly(A)^+^ RNA-FISH. Scale bar = 20 μm. (**F**) Quantification of proportion of cytoplasmic RNA granule^+^ cells in (E). Dunnett’s multiple-comparison test, ***P < 0.001. Error bars indicate mean ± SE. N = 3 independent experiments.

In addition to a decrease in the percentage of cells containing mitoRGs, the size of the granules also decreased from 8–12 h after CCCP treatment (**Fig. 1A–D**). These results suggest that prolonged mitochondrial stress eventually leads to mitoRG disassembly. Moreover, CCCP-induced mitoRG formation was not observed in HeLa cells, in which the endogenous level of Parkin is known to be insufficient to induce mitophagy (Narendra *et al*, 2008) (**Fig. S1C**). Overall, mitoRGs appear to be formed transiently after the induction of PINK1/Parkin mitophagy and disassembled after prolonged mitochondrial stress.

To determine if PGAM5 affected the formation of mitoRGs during PINK1/Parkin mitophagy, we performed poly(A)^+^ RNA-FISH in PGAM5-deficient Parkin-HeLa cells (PGAM5 knockout (KO) Parkin-HeLa cells), as established previously (Baba *et al*, 2021). CCCP-induced formation of mitoRGs was markedly suppressed in PGAM5 KO cells compared with wild-type cells (**Fig. 1E, F**), suggesting that PGAM5 is required for mitoRG formation.

### mitoRGs differ from SGs in terms of components and formation mechanisms

Next, we compared the features of mitoRGs with those of stress granules (SGs), which are well-characterized cytoplasmic mRNA granules formed in response to various cellular stressors (Marcelo *et al*, 2021). Simultaneous immunofluorescence and poly(A)^+^ RNA-FISH analyses revealed that the RNA-binding protein G3BP1, which acts as a key nucleator of SGs, co-localized with mitoRGs in Parkin-HeLa cells treated with CCCP (**Fig. 2A, B**) or OA (**Fig. S2A**). In line with the lack of mitoRGs in HeLa cells (**Fig. S1C**), no obvious G3BP1 granules were observed in cells treated with CCCP (**Fig. S2B**). These results suggest that G3BP1 is a component of mitoRGs and thus serves as a marker for mitoRGs. In fact, G3BP1 immunofluorescence could assess the difference in the extent of CCCP-induced mitoRG formation between wild-type and PGAM5 KO Parkin-Hela cells (**Fig. 2C, D**) and between Parkin-HeLa cells in which PGAM5 expression was suppressed with small interfering RNAs (siRNAs) and those transfected with control siRNAs (**Fig. S2C, D**). In contrast, sodium arsenite-induced assembly of SGs, which was also assessed using G3BP1 immunofluorescence, was not attenuated in PGAM5 KO Parkin-HeLa cells (**Fig. 2E, F**), suggesting that the mechanisms by which cytoplasmic mRNA granules are assembled differ between mitoRGs and SGs.

**Figure 2.**
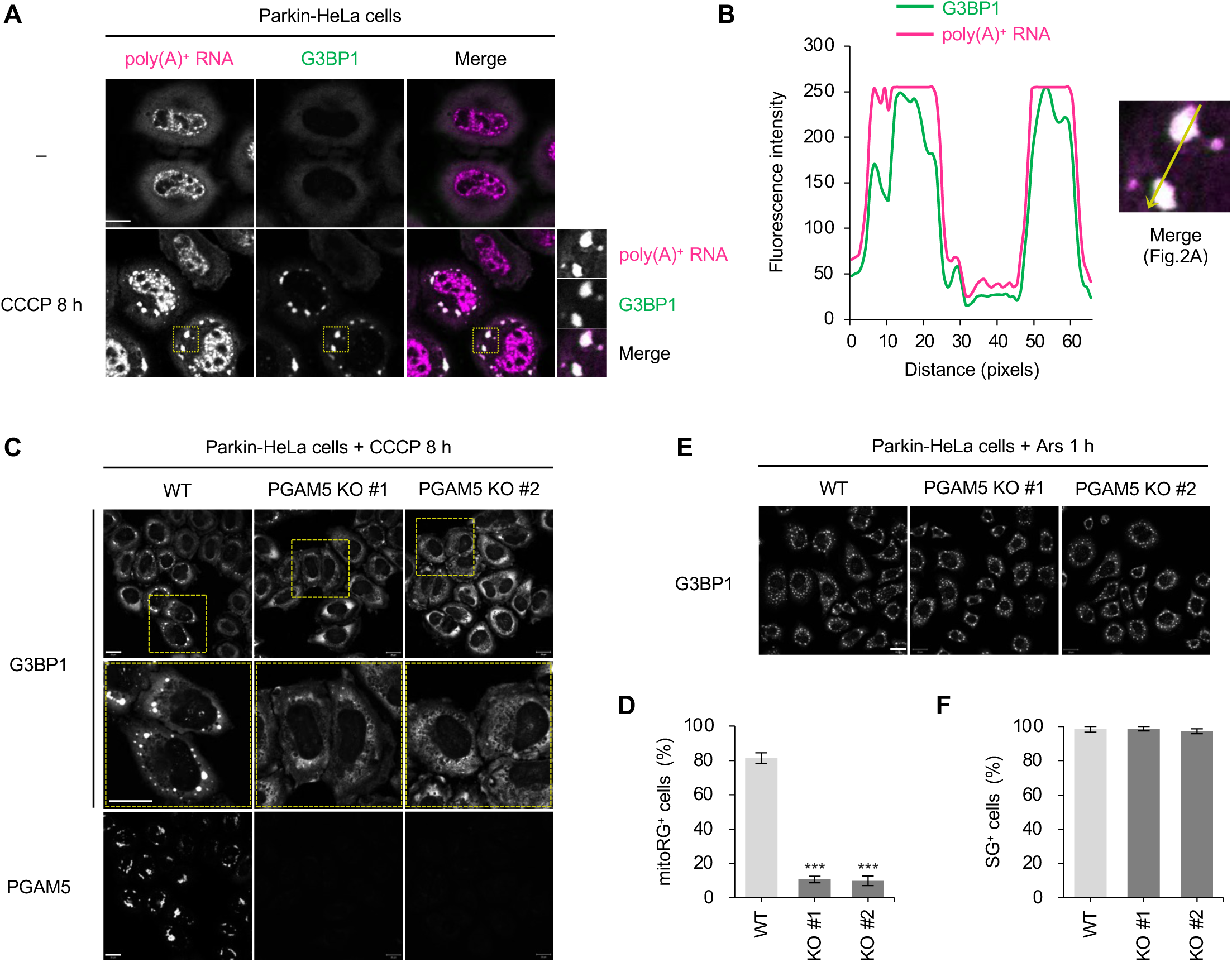
PGAM5 is required for the assembly of mitoRGs but not for that of SGs. (**A**) Representative poly(A)^+^ RNA-FISH and immunofluorescence results for G3BP1 in Parkin-HeLa cells treated with 10 μM CCCP for 8 h. Scale bar = 20 μm. Magnified images of areas outlined by yellow rectangles in images of CCCP-treated cells are shown in lower-right panels. (**B**) Line profiles of fluorescence intensities of G3BP1 (green) and poly(A)^+^ RNA (magenta) in CCCP-treated cells (left graph), generated along the plane indicated by the yellow arrow in the inset of the merged image in (A) (right panel). (**C, E**) Parkin-HeLa cells (WT) and two lines of PGAM5 KO Parkin-HeLa cells (PGAM5 KO #1 and #2) were treated with 10 μM CCCP for 8 h (C) or 0.25 mM sodium arsenite (Ars) for 1 h (E) and subjected to immunofluorescence analysis for G3BP1 (C, E) and PGAM5 (C). Magnified images of areas outlined by yellow rectangles in G3BP1 images are shown in middle panels. Scale bar = 20 μm (**D, F**). Quantification of proportions of mitoRG^+^ cells in (C) and SG^+^ cells in (D). Dunnett’s multiple-comparison test, ***P < 0.001. Error bars indicate mean ± SE. N = 3 independent experiments.

We further examined whether the other components of SGs, the eukaryotic translation initiation factor eIF4G, the RNA-binding protein TIA1, and the 40S ribosomal subunits RPS3 and RPS6, are involved in both SGs and mitoRGs. Although both eIF4G and TIA1 were incorporated into arsenite-induced SGs and CCCP-induced mitoRGs, RPS3 and RPS6 were present in SGs but not in mitoRGs (**Fig. 3A, B**). The inclusion of RPS6 in SGs was also confirmed in energy deficiency-induced SGs (eSG), which were noncanonically induced under severe ATP depletion (Wang *et al*, 2022) (**Fig. S3A**), suggesting that mitoRGs are different from SGs, irrespective of their inducers, in terms of the inclusion of 40S ribosomal subunits. A previous study reported that G3BP1 and 40S ribosomal subunits interacted with each other and coordinately mediated SG condensation as key nucleators (Kedersha *et al*, 2016). The difference in the involvement of the 40S ribosomal subunits between mitoRGs and SGs may thus reflect the differences in their assembly mechanisms.

**Figure 3.**
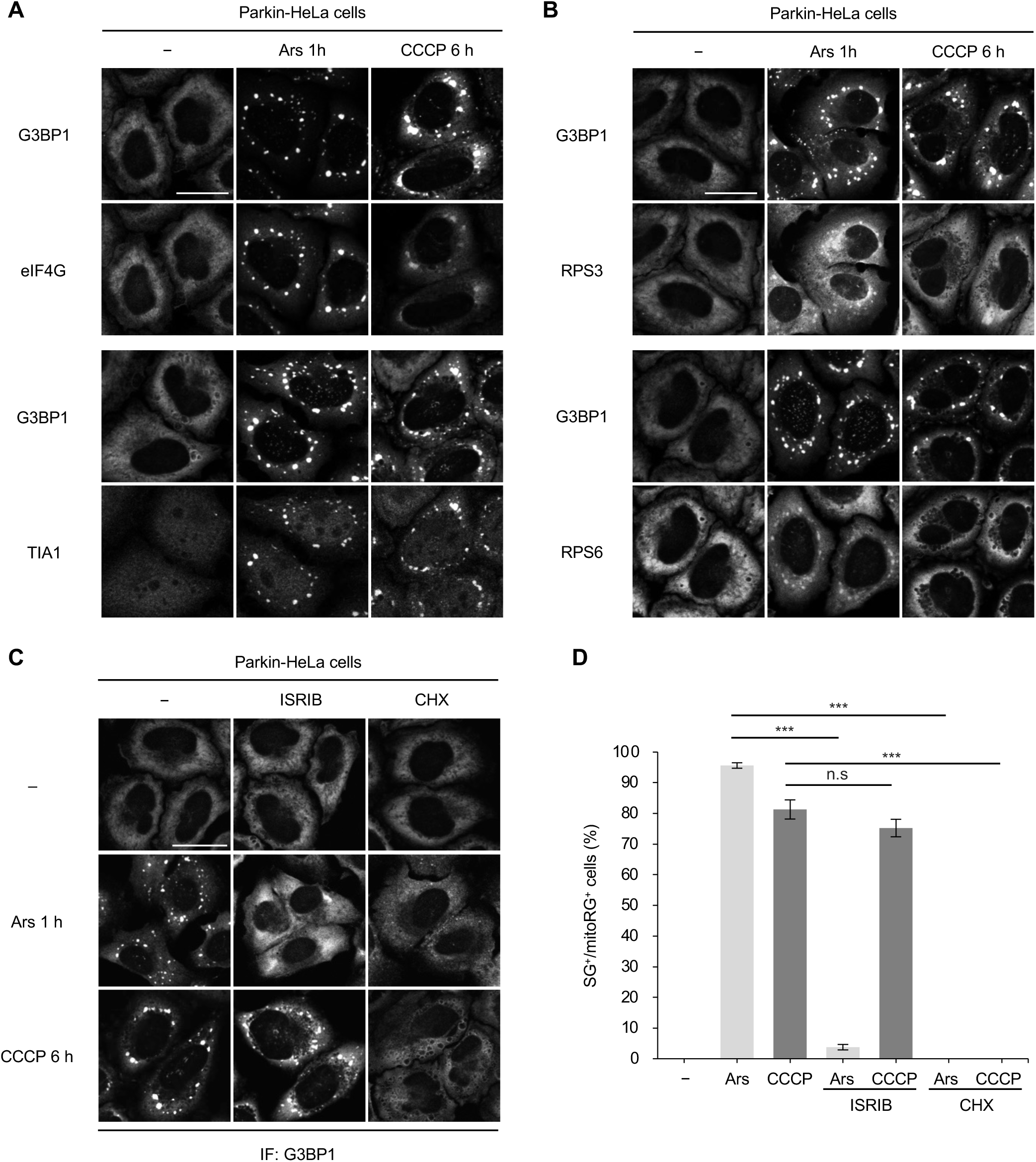
mitoRGs differ from SGs in terms of components and formation mechanisms. (**A, B**) Representative dual immunofluorescence results using G3BP1 antibody and either eIF4G (A), TIA1 (A), RPS3 (B), or RPS6 (B) antibody in Parkin-HeLa cells treated with 0.25 mM sodium arsenite (Ars) for 1 h or 10 μM CCCP for 6 h. Scale bar = 20 μm. (**C**) Parkin-HeLa cells pretreated with 0.5 μM ISRIB or 10 μg/ml cycloheximide (CHX) were treated with 0.25 mM Ars for 1 h or 10 μM CCCP for 6 h and subjected to immunofluorescence (IF) for G3BP1. Scale bar = 20 μm. (**D**) Quantification of proportions of SG^+^ and mitoRG^+^ cells in (C). Tukey’s multiple-comparison test, ***P < 0.001. Error bars indicate mean ± SE. N = 3 independent experiments.

The assembly of SGs requires free mRNAs released from ribosomes following a global translation arrest (Bounedjah *et al*, 2014). We therefore examined whether and how translation arrest occurs prior to mitoRG assembly. To quantify global translation levels, we performed a puromycin incorporation assay (Schmidt *et al*, 2009). In this assay, cultured cells were pulsed with puromycin, which was incorporated into nascent peptides, and the puromycylated peptides were subsequently detected by immunoblotting with puromycin antibody (**Fig. S3B**). Broad smear bands were detected under steady-state conditions, whereas arsenite treatment caused a marked reduction in band intensity, reflecting robust suppression of global translation (**Fig. S3C**). Similarly, CCCP treatment significantly reduced band intensity, indicating that global translation arrest occurs before mitoRG formation.

Translational arrest during SG assembly is typically mediated by eIF2α phosphorylation (P-eIF2α), which inhibits translation initiation (McCormick & Khaperskyy, 2017). However, no detectable eIF2α phosphorylation was observed during CCCP- or OA-induced mitophagy (**Fig. S3E, F**). Furthermore, integrated stress response inhibitor (ISRIB), a small molecule that reverses eIF2α-dependent translation arrest (Sidrauski *et al*, 2013), strongly suppressed arsenite-induced SG formation but had almost no effect on mitoRG assembly (**Fig. 3C, D**), suggesting that mitoRGs are assembled through an eIF2α-independent mechanism of translation inhibition. To determine if mitoRG formation required polysome disassembly, as for SGs, we treated cells with arsenite or CCCP in the presence of cycloheximide, which inhibits the polysome disassembly and consequently blocks subsequent translation elongation. Cycloheximide significantly inhibited both SG and mitoRG formation (**Fig. 3C, D**), suggesting that mitoRG formation is dependent on polysome disassembly. Overall, these results demonstrate that mitoRGs are assembled via a unique pathway that depends on polysome disassembly, but not on eIF2α.

### PGAM5 facilitates mitoRG formation through intramolecular cleavage and phosphatase activity

We next investigated the role of PGAM5 in mitoRG formation. We have previously found that PGAM5 is a protein phosphatase that requires His105 in the PGAM domain for its activity (Takeda *et al*, 2009) and Ser24 for intramolecular cleavage upon mitochondrial stress (Sugawara *et al*, 2020). Exogenous expression of wild-type PGAM5, but not the phosphatase-dead mutant PGAM5 H105A, rescued mitoRG formation in PGAM5 KO cells (**Fig. 4A, B**), suggesting that mitoRG formation depends on the phosphatase activity of PGAM5. Exogenous expression of the cleavage-resistant mutant S24W (Sugawara *et al*, 2020) also failed to rescue mitoRG formation. These observations led us to hypothesize that cleaved PGAM5, released from mitochondria during mitophagy, facilitates mitoRG formation. To test this hypothesis, we treated cells with the proteasome inhibitor MG132 to suppress the proteasome-dependent rupture of the mitochondrial outer membrane, which is required for the release of cleaved PGAM5 from mitochondria during mitophagy (Yamaguchi *et al*, 2019). As expected, mitoRG formation was strongly inhibited under these conditions (**Fig. 4C**). Moreover, a portion of PGAM5, but not the mitochondrial matrix protein ATP5B or cytochrome c, the latter of which is released from mitochondria during mitophagy, co-localized with mitoRGs. (**Fig. 4D–F**). Collectively, these results suggest that, upon release from mitochondria, cleaved PGAM5 translocates to mitoRGs and promotes their assembly in a phosphatase activity-dependent manner.

**Figure 4.**
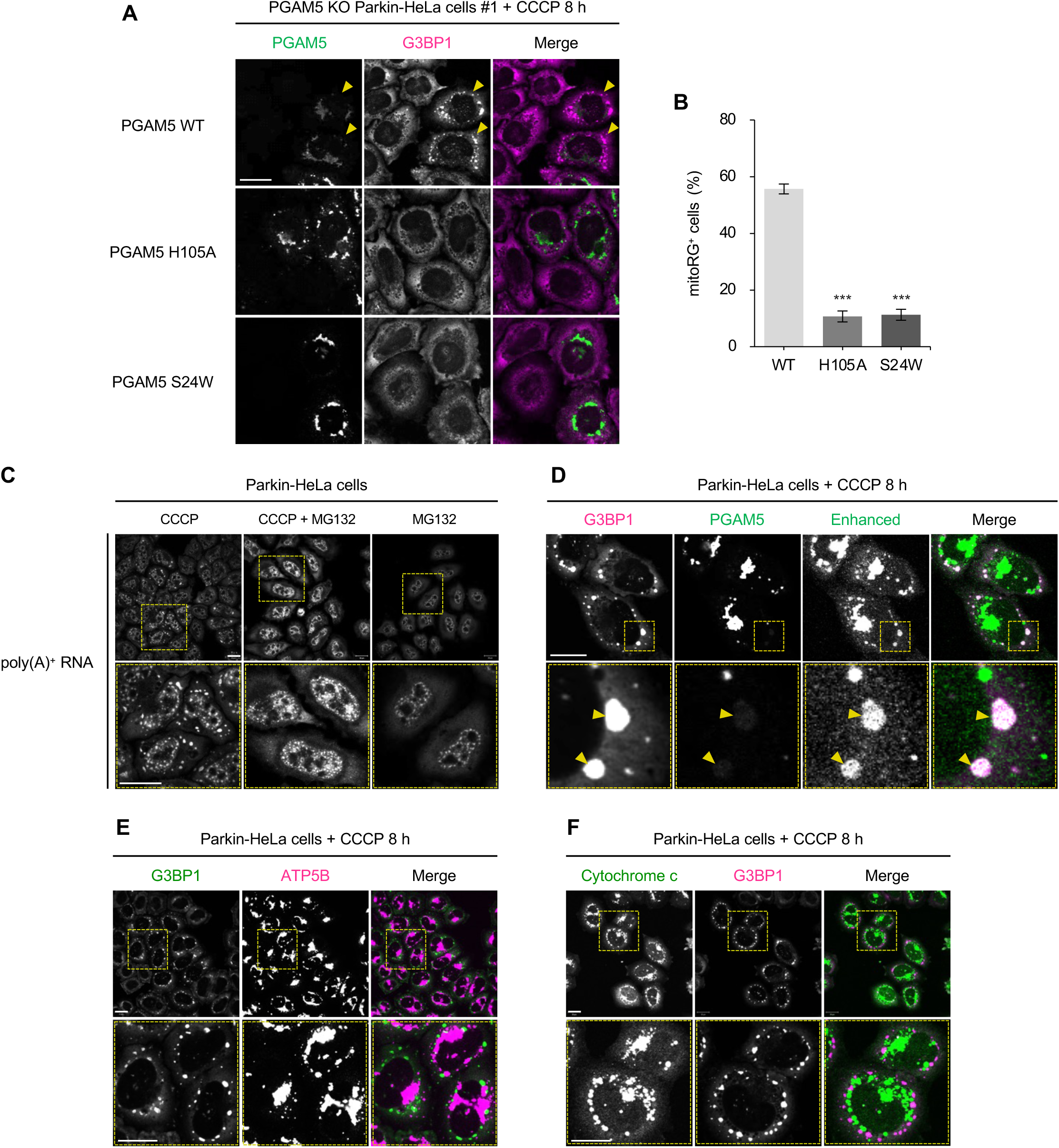
PGAM5 facilitates mitoRG formation through intramolecular cleavage and phosphatase activity. (**A**) PGAM5 KO Parkin-HeLa cells (#1) exogenously expressing PGAM5 wild-type (WT), H105A, and S24W were treated with 10 μM CCCP for 8 h and subjected to immunofluorescence analysis for PGAM5 and G3BP1. Scale bar = 20 μm. Yellow arrowheads indicate the cells expressing PGAM5 WT. (**B**) Quantification of proportion of mitoRG^+^ cells in (A). Dunnett’s multiple-comparison test, ***P < 0.001. Error bars indicate mean ± SE. N = 3 independent experiments. (**C**) Representative results of poly(A)^+^ RNA-FISH in Parkin-HeLa cells treated with 5 μM MG132, followed by the 8-h treatment with 10 μM CCCP for 8 h (upper panels). Magnified images of areas outlined by yellow rectangles in upper panels are shown in lower panels. Scale bar = 20 μm. (**D**) Representative immunofluorescence results for G3BP1 and PGAM5 in Parkin-HeLa cells treated with 10 μM CCCP for 8 h. Scale bar = 20 μm. Magnified images of areas outlined by yellow rectangles in upper panels are shown in lower panels. Yellow arrowheads indicate the co-localization of PGAM5 and G3BP1. (**E, F**) Representative dual immunofluorescence results using G3BP1 antibody and either ATP5B (E) or cytochrome c (F) antibody in Parkin-HeLa cells treated with 10 μM CCCP for 8 h. Scale bar = 20 μm.

### G3BP1 depletion inhibits mitoRG formation and promotes apoptosis during PINK1/Parkin mitophagy

Next, we investigated the significance of mitoRG formation in the cellular response to mitochondrial stress that induces PINK1/Parkin mitophagy. CCCP-induced mitoRG formation was inhibited in Parkin-HeLa cells in which G3BP1 was knocked down using siRNAs (**Fig. 5A–C**), suggesting that G3BP1 acts as a key nucleator of mitoRGs, as for SGs. In this experiment, we noticed that G3BP1 knockdown cells were more vulnerable than control cells (**Fig. 5D**). Previous studies have reported that mitochondrial depolarization in the presence of Parkin initially activates mitophagy as a cytoprotective response, and that prolonged depolarization, which induces sustained mitochondrial stress, shifts the cellular outcome toward apoptosis (Quarato *et al*, 2023; Carroll *et al*, 2014). Consistent with these findings, Parkin-HeLa cells exhibited apoptotic features after 12-h treatment with CCCP, as evidenced by significantly increased SYTOX Green-positive dead cells, and this effect was suppressed by the pan-caspase inhibitor Q-VD (**Fig. 5E**). However, immunofluorescence analysis of cleaved PARP, as a representative marker of apoptosis, detected apoptotic cells in G3BP1 knockdown cells but not in control cells 8 h after CCCP treatment (**Fig. 5F, G**). These results suggest that mitoRG formation plays a protective role by suppressing apoptosis induced by sustained mitochondrial stress.

**Figure 5.**
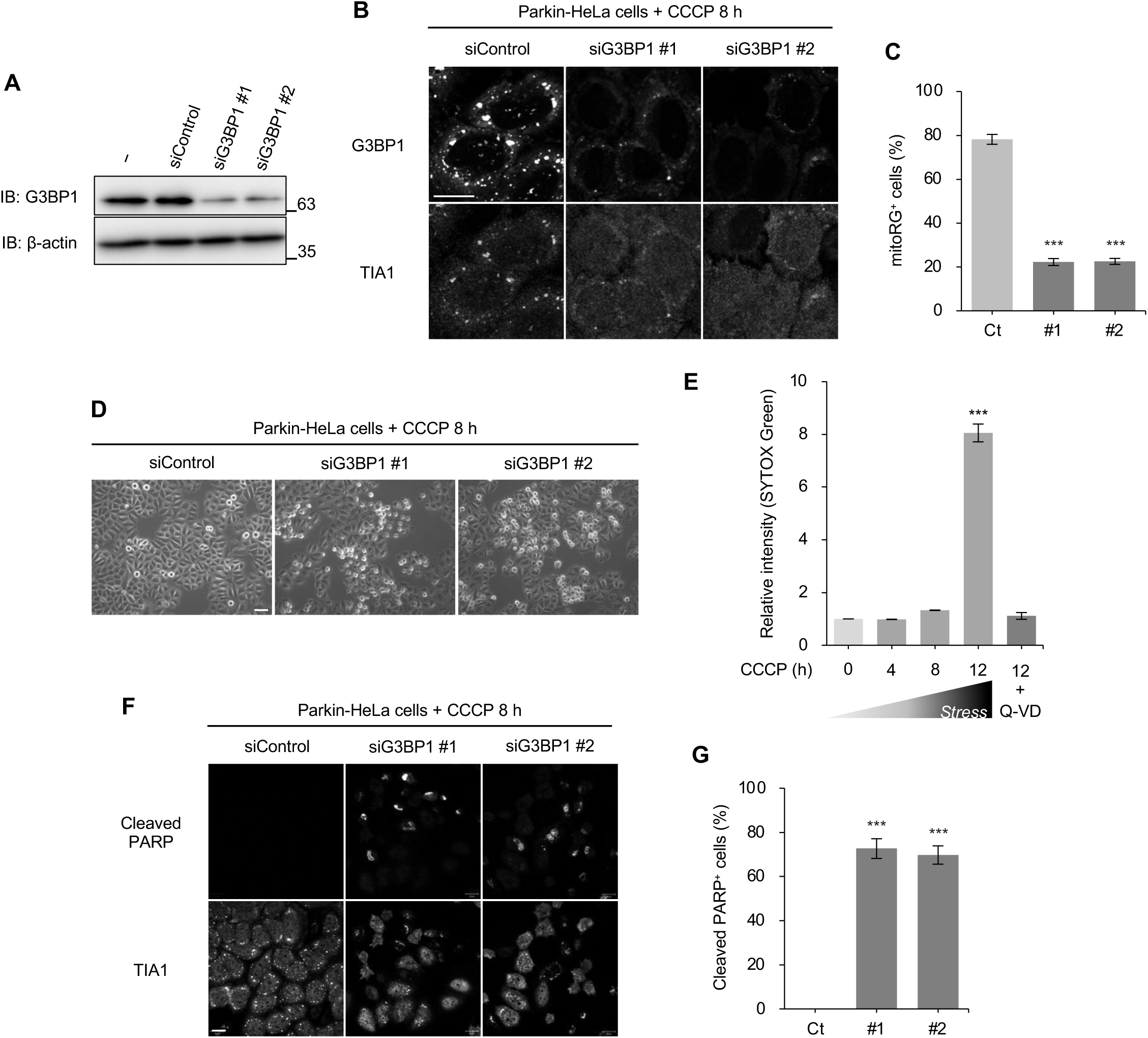
G3BP1 depletion inhibits mitoRG formation and promotes apoptosis during PINK1/Parkin mitophagy. (**A**) Parkin-HeLa cells transfected with siRNAs targeting G3BP1 (siG3BP1 #1 and #2) or control siRNA (siControl) were subjected to immunoblotting (IB) with G3BP1 and β-actin antibodies. (**B**) Parkin-HeLa cells transfected with siG3BP1 #1, #2, or siControl were treated with 10 μM CCCP for 8 h and subjected to immunofluorescence analysis for G3BP1 and TIA1. Scale bar = 20 μm. (**C**) Quantification of proportion of mitoRG^+^ cells in (B). Dunnett’s multiple-comparison test, ***P < 0.001. Error bars indicate mean ± SE. N = 3 independent experiments. (**D**) Representative phase-contrast images of Parkin-HeLa cells transfected with siG3BP1 #1, #2, or siControl, followed by treatment with 10 μM CCCP for 8 h. Scale bar = 50 μm. (**E**) Parkin-HeLa cells were treated with 10 μM CCCP for 4, 8, and 12 h and with 10 μM CCCP for 12 h in the presence of 10 μM Q-VD, followed by staining with SYTOX Green. Graph showing relative fluorescence intensity of SYTOX Green compared with control (CCCP 0 h). Dunnett’s multiple-comparison test, ***P < 0.001. Error bars indicate mean ± SE. N = 3 independent experiments. (**F**) Parkin-HeLa cells transfected with siG3BP1 #1, #2, or siControl were treated with 10 μM CCCP for 8 h and subjected to immunofluorescence analysis for cleaved PARP and TIA1. Scale bar = 20 μm. (**G**) Quantification of proportion of cleaved PARP^+^ cells in (F). Dunnett’s multiple-comparison test, ***P < 0.001. Error bars indicate mean ± SE. N = 3 independent experiments.

### Impaired mitoRG formation promotes apoptosis in a PGAM5-dependent manner during PINK1/Parkin mitophagy

As the impairment of mitoRGs by G3BP1 knockdown sensitizes cells to apoptosis, it is conceivable that apoptosis is also promoted in PGAM5 KO cells, which exhibit impaired mitoRG formation. Contrary to this expectation, apoptosis was suppressed, rather than promoted, in PGAM5 KO cells compared with wild-type cells, as demonstrated by phase-contrast imaging (**Fig. 6A**) and terminal deoxynucleotidyl transferase (TdT)-mediated dUTP nick-end labeling (TUNEL) staining (**Fig. 6B**). Consistent with these results, CCCP-induced increases in apoptosis markers, such as cleaved caspase-9, cleaved caspase-3, and cleaved PARP, were more attenuated in PGAM5 KO cells than in wild-type cells (**Fig. 6C, D**). These results led us to hypothesize that the impairment of mitoRG formation promotes apoptosis depending on PGAM5. To test this hypothesis, we knocked down G3BP1 using siRNA in PGAM5 KO cells, treated the cells with CCCP, and subsequently assessed apoptosis by immunostaining for cleaved PARP. As expected, the increase in apoptotic cells following G3BP1 knockdown was strongly suppressed in PGAM5 KO cells (**Fig. 6E, F**). Collectively, these results suggest that the impairment of mitoRG formation promotes apoptosis in a PGAM5-dependent manner during PINK1/Parkin mitophagy.

**Figure 6.**
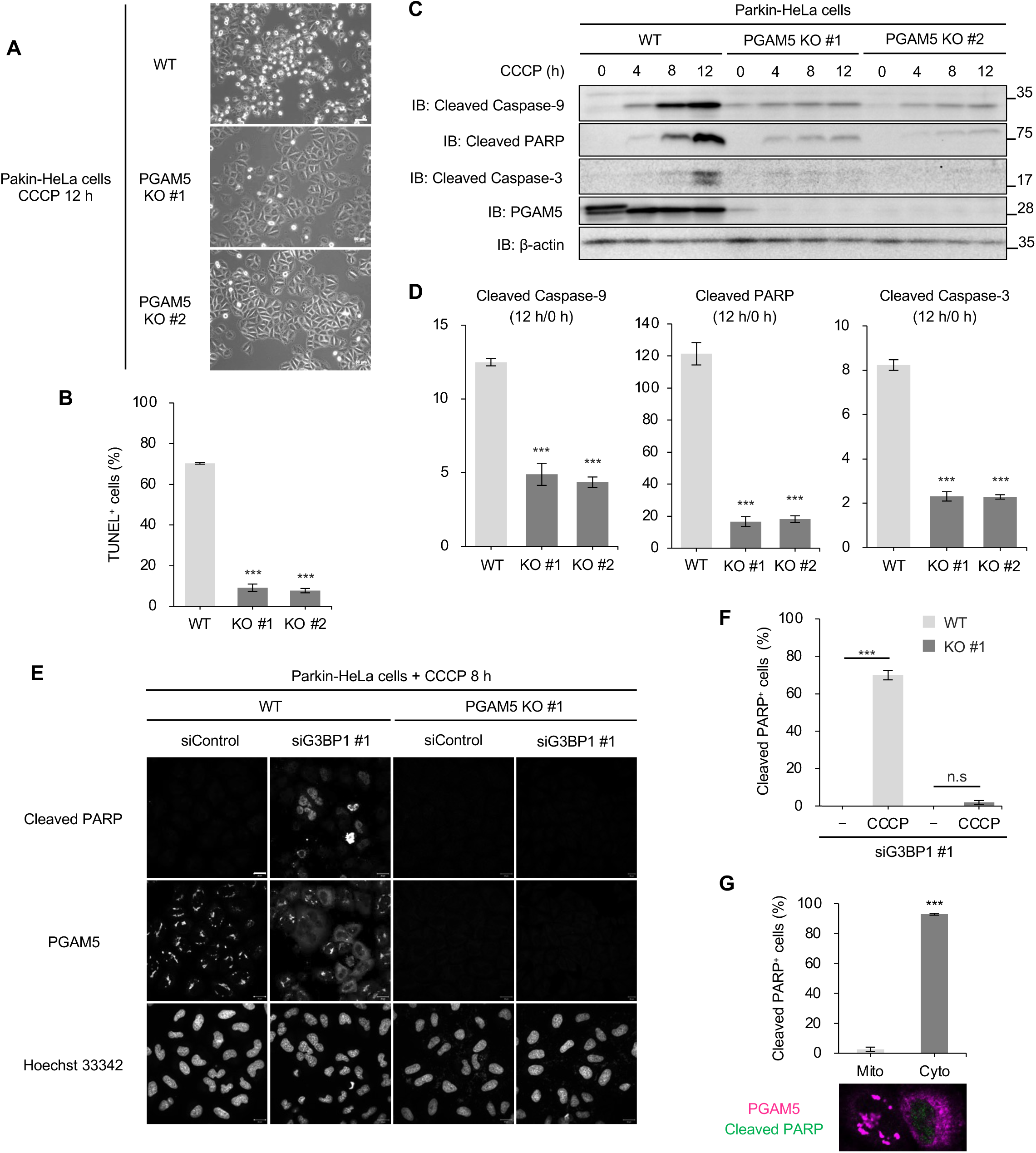
Impaired mitoRG formation promotes apoptosis in a PGAM5-dependent manner during PINK1/Parkin mitophagy. (**A**) Representative phase-contrast images of Parkin-HeLa (WT) and PGAM5 KO Parkin-HeLa cells (PGAM5 KO #1 and KO #2) treated with 10 μM CCCP for 12 h. Scale bar = 50 μm. (**B**) TUNEL assay in Parkin-HeLa cells (WT, PGAM5 KO #1, and #2) treated with 10 μM CCCP for 12 h. The proportion of TUNEL^+^ cells was quantified. Dunnett’s multiple-comparison test, ***P < 0.001. Error bars indicate mean ± SE. N = 3 independent experiments. (**C**) Parkin-HeLa cells (WT, PGAM5 KO #1, and #2) were treated with 10 μM CCCP for 4, 8, and 12 h and subjected to immunoblotting (IB) with the indicated antibodies. (**D**) Fold changes in apoptosis markers (12 h versus 0 h) in (C). Dunnett’s multiple-comparison test, ***P < 0.001. Error bars indicate mean ± SE. N = 3 independent experiments. (**E**) Parkin-HeLa cells (WT and PGAM5 KO #1) transfected with siRNA targeting G3BP1 (siG3BP1 #1) or control siRNA (siControl) were treated with 10 μM CCCP for 8 h and subjected to immunofluorescence analysis for cleaved PARP and PGAM5, followed by counterstaining with Hoechst 33342. Scale bar = 20 μm. (**F**) The proportion of cleaved PARP^+^ cells in (E) was quantified. Tukey’s multiple-comparison test, ***P < 0.001. Error bars indicate mean ± SE. N = 3 independent experiments. (**G**) Relationship between subcellular localization of PGAM5 and proportion of cleaved PARP^+^ cells in Parkin-HeLa cells transfected with siG3BP1 #1 and treated with 10 μM CCCP for 8 h in (E). *Mito*, cells in which PGAM5 is retained in mitochondria; *Cyto*, cells in which PGAM5 is diffusely distributed. Student’s t-test, ***P < 0.001. Error bars indicate mean ± SE. N = 3 independent experiments. Representative images of the two distribution patterns of PGAM5 are shown below the graph.

In addition, G3BP1 knockdown caused cytosolic diffusion of PGAM5 in most wild-type cells (middle, second panel from the left in **Fig. 6E**). Cleaved PARP-positive apoptotic cells were induced in cells exhibiting cytosolic PGAM5 localization but not in those exhibiting mitochondrial PGAM5 localization (**Fig. 6G**). These results suggest that cytosolic diffusion of PGAM5 is a critical factor linking impaired mitoRG formation to the execution of apoptosis under mitochondrial stress.

### mitoRGs transiently sequester PGAM5 to inhibit its cytosolic diffusion and pro-apoptotic function during PINK1/Parkin mitophagy

PGAM5 released from mitochondria translocated to mitoRGs (**Fig. 4D**), and was diffusely distributed throughout the cytosol when mitoRGs failed to form, such as after knockdown of G3BP1 (**Fig. 6E**). These results suggest that mitoRGs sequester PGAM5 from the cytosol, thereby preventing its uncontrolled cytosolic accumulation and the subsequent execution of apoptosis under mitochondrial stress. We examined this hypothesis by time-course immunostaining for PGAM5 and G3BP1 in Parkin-HeLa cells treated with CCCP. Consistent with the poly(A)^+^ RNA-FISH results (**Fig. 1A, B**), mitoRGs progressively diminished in terms of both size and number at 8–12 h after CCCP treatment (**Fig. 7A**). This temporal window of mitoRG shrinkage and dissolution coincided with the onset of PGAM5 diffusion into the cytosol during CCCP-induced (**Fig. 7A**) and OA-induced mitophagy (**Fig. S4A**). To examine the distribution of PGAM5 more precisely, we treated cells with CCCP in the presence of Q-VD to inhibit PGAM5-dependent apoptosis, and found that impaired mitoRG formation upon G3BP1 knockdown resulted in the diffuse cytosolic localization of PGAM5 in most cells (**Fig. 7B, C**).

**Figure 7.**
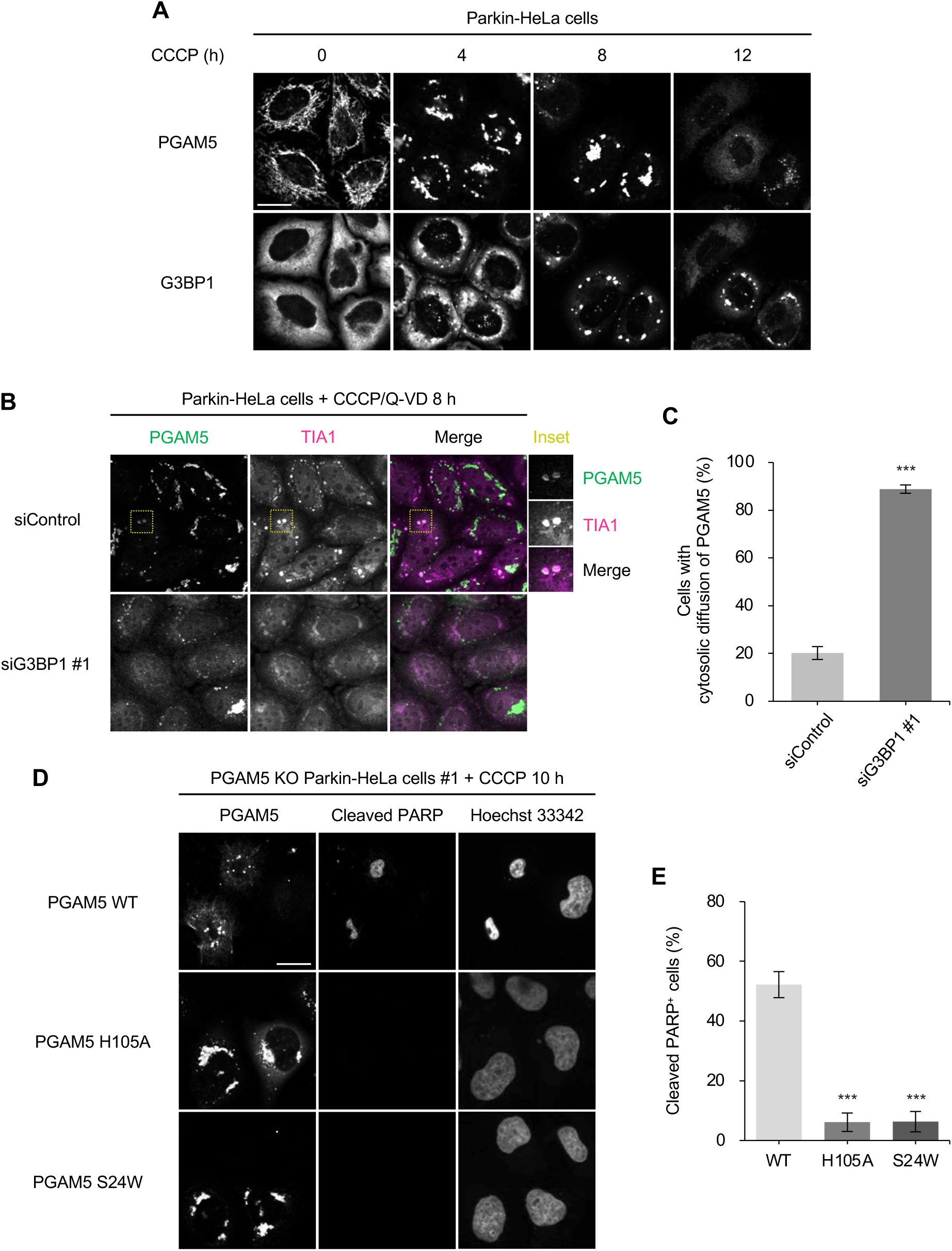
mitoRGs transiently sequester PGAM5 to inhibit its cytosolic diffusion and pro-apoptotic function during PINK1/Parkin mitophagy. (**A**) Representative immunofluorescence results for PGAM5 and G3BP1 in Parkin-HeLa cells treated with 10 μM CCCP for 4, 8, and 12 h. Scale bar = 20 μm. (**B**) Parkin-HeLa cells transfected with siRNA targeting G3BP1 (siG3BP1) or control siRNA (siControl) were treated with 10 μM CCCP and 20 μM Q-VD for 8 h and subjected to immunofluorescence analysis for PGAM5 and TIA1. Scale bar = 20 μm. Magnified images of areas outlined by yellow rectangles in upper large panels are shown in upper-right small panels. (**C**) The proportion of cells that showed cytosolic diffusion of PGAM5 in (B) was quantified. Student’s t-test, ***P < 0.001. Error bars indicate mean ± SE. N = 3 independent experiments. (**D**) Representative immunofluorescence results for cleaved PARP and PGAM5 in PGAM5 KO Parkin-HeLa cells (#1) exogenously expressing PGAM5 wild-type (WT), H105A, or S24W treated with 10 μM CCCP for 10 h, followed by counterstaining with Hoechst 33342. Scale bar = 20 μm. (**E**) The proportion of cleaved PARP^+^ cells in (D) was quantified. Dunnett’s multiple-comparison test, ***P < 0.001. Error bars indicate mean ± SE. N = 3 independent experiments.

These results reinforce the idea that mitoRGs may be required to constrain mitochondria-derived PGAM5 within a defined subcellular compartment and prevent its widespread cytosolic diffusion. Thus, mitoRG disassembly may enable the release of PGAM5 into the cytosolic compartment. Consistent with this idea, a subset of Parkin-HeLa cells that exhibited diffuse cytosolic localization of PGAM5 was simultaneously positive for cleaved PARP at 12 h after CCCP treatment, when mitoRGs strongly decreased (**Fig. S4B**), suggesting that PGAM5 released into the cytosol upon mitoRG disassembly induces apoptosis. This was supported by the rescue of apoptosis in PGAM5 KO cells 10 h after CCCP treatment by the exogenous expression of wild-type PGAM5, but not by expression of the phosphatase-dead mutant H105A or the cleavage-resistant mutant S24W (**Fig. 7D, E**). Overall, following mitochondrial stress-induced cleavage and release from mitochondria, PGAM5 appears to be initially sequestered within mitoRGs but may subsequently be released into the cytosol, triggering apoptosis.

## Discussion

In this study, we identified a previously unrecognized class of cytoplasmic mRNA granules that assemble transiently during PINK1/Parkin mitophagy, which we named mitoRGs. The results demonstrated that mitoRGs are similar to but distinct from SGs and play a critical role in cell fate decisions under mitochondrial stress by spatiotemporally regulating the localization and pro-apoptotic activity of the mitochondrial protein phosphatase PGAM5.

Although SGs assemble in response to diverse stress conditions, including oxidative stress, heat shock, and ER stress, mitoRGs differ from SGs in several fundamental respects. SG assembly typically depends on translational arrest mediated by eIF2α phosphorylation downstream of stress-activated kinases, such as PERK, GCN2, PKR, and HRI (Pakos-Zebrucka *et al*, 2016). In contrast, mitoRG formation occurs independently of eIF2α phosphorylation and is insensitive to ISRIB, a small molecule that restores translation under conditions of integrated stress response (Sidrauski *et al*, 2013), indicating that mitoRGs are not under the control of canonical integrated stress response signaling. Canonical and non-canonical SGs, including energy stress-induced SGs (eSGs), contain stalled translation preinitiation complexes and 40S ribosomal subunits that serve as critical nucleation scaffolds (Wang *et al*, 2022; Kedersha *et al*, 2016). In this regard, the absence of 40S ribosomal proteins from mitoRGs indicates distinct granule composition and assembly mechanisms. In addition, SG formation can be triggered by a wide range of cellular stresses, whereas mitoRG assembly is induced by mitochondrial dysfunction associated with PINK1/Parkin mitophagy, a pathway that marks damaged mitochondria for autophagic degradation (Narendra & Youle, 2024). Collectively, these findings strongly suggest that mitoRGs comprise a distinct class of mRNA granules that are mechanistically different from SGs. Despite these differences, mitoRGs share several features with SGs, including dependence on polysome disassembly and incorporation of the RNA-binding protein G3BP1, which is a central SG nucleator (Yang *et al*, 2020). These partially overlapping aspects suggest that mitoRGs repurpose the core mRNA condensation machinery to generate a functionally specialized granule that integrates mitochondrial stress-specific signals, consistent with the emerging view that the modular reuse of RNA granule components underlies the diversity of ribonucleoprotein condensates (Mittag & Parker, 2018).

Our results further identified the mitochondrial protein phosphatase PGAM5 as a key molecular switch linking mitochondrial damage to RNA granule dynamics and apoptotic signaling. PGAM5 is an atypical Ser/Thr protein phosphatase localized to the inner mitochondrial membrane (Sekine *et al*, 2012; Takeda *et al*, 2009), which has been implicated in multiple mitochondrial stress responses, including the regulation of mitochondrial dynamics and apoptosis (Nag *et al*, 2025; Ma *et al*, 2020; Nag *et al*, 2023). In response to mitochondrial depolarization, PGAM5 is cleaved by PARL or OMA1 and released into the cytosol, where it serves as a signaling intermediate that induces various cellular responses, such as caspase activation and apoptosis (Saita *et al*, 2017; Zhuang *et al*, 2013). Consistent with the molecular behavior of PGAM5, we found a correlation between the cytosolic diffusion of PGAM5 and the onset of apoptosis, both of which occurred during prolonged mitochondrial stress but not as early events. This suggests that PGAM5 does not necessarily exert its pro-apoptotic function immediately after its release from mitochondria into the cytosol and that a certain mechanism restrains PGAM5 from exerting its pro-apoptotic activity. Regarding this mechanism, PGAM5 appears to be initially sequestered within mitoRGs, thereby restricting its diffuse cytosolic distribution. The spatial sequestration of signaling molecules by biomolecular condensates has emerged as a powerful mechanism for modulating the amplitude and timing of signaling (Fujikawa *et al*, 2023; Shin & Brangwynne, 2017). Thus, mitoRGs function as a buffering platform that temporally regulates PGAM5-driven apoptotic signaling. This was supported by the data showing that impairment of mitoRG assembly through G3BP1 depletion resulted in the unconfined cytosolic distribution of PGAM5 and PGAM5-dependent promotion of apoptosis during mitophagy. These findings suggest that mitoRGs extend the temporal window during which cells are allowed to restore mitochondrial homeostasis to survive.

The current study also demonstrated the capacity of PGAM5 to engage RNA regulatory pathways across multiple cellular compartments in a temporally ordered manner. We previously showed that PGAM5 released from mitochondria translocated to the nucleus during PINK1/Parkin mitophagy, where it dephosphorylated nuclear proteins involved in mRNA metabolism, including those implicated in mRNA processing, export, and stability (Baba *et al*, 2021). Perturbations in mRNA metabolism are closely linked to cell fate decisions, and defects in mRNA processing or export can induce apoptosis (Bowling *et al*, 2021; Moore *et al*, 2010; Chappaz *et al*, 2020). Our temporal analyses indicated that the nuclear translocation of PGAM5 and dephosphorylation of nuclear mRNA metabolic regulators occurred predominantly at later stages of mitochondrial stress, coinciding with mitoRG shrinkage or dissolution and the onset of apoptosis. Furthermore, cells in which PGAM5 dephosphorylated nuclear factors were coincidentally positive for cleaved PARP, suggesting that the modulation of nuclear mRNA metabolism by PGAM5 is linked to the response to prolonged mitochondrial stress rather than to the earlier cytoprotective phase of mitophagy (**Fig. S5**). Despite a lack of evidence showing that PGAM5-mediated dephosphorylation of nuclear proteins is directly coupled to mitoRG dynamics, mitoRGs appear to function primarily to spatially confine mitochondria-derived PGAM5 within the cytoplasm during the early phase of mitophagy, thereby delaying the access of PGAM5 to cytosolic pro-apoptotic effectors and nuclear substrates. Under conditions of prolonged mitochondrial stress, mitoRG disassembly permits widespread cytosolic redistribution of PGAM5 and facilitates its subsequent nuclear translocation. In this context, PGAM5-mediated dephosphorylation of nuclear proteins involved in mRNA metabolism may contribute to apoptosis by modulating RNA processing, export, or surveillance pathways, reinforcing a transcriptional and post-transcriptional state incompatible with cell survival.

The identification of mitoRGs has revealed an unanticipated role for RNA granules in mitochondrial quality control and cell fate regulation. Given the central involvements of PINK1/Parkin signaling and PGAM5 in neurodegenerative diseases, particularly Parkinson’s disease (Pickrell & Youle, 2015; Valente *et al*, 2004; Lu *et al*, 2014), the dysregulation of mitoRG dynamics may contribute to the pathological outcomes associated with mitochondrial dysfunction. However, several important questions remain, including the identity of the mRNA populations associated with mitoRGs, the molecular targets of PGAM5 within these granules, and the mechanisms that trigger mitoRG disassembly under prolonged stress. Addressing these issues will further elucidate how mitochondrial stress is integrated into post-transcriptional regulatory networks to control cell fate.

In summary, our study identified mitoRGs as a novel mRNA-based regulatory layer that coordinates mitochondrial quality control with apoptosis through the spatiotemporal regulation of PGAM5. This study expands the conceptual landscape of stress-responsive mRNA granules and provides new insights into how cells balance survival and death in response to mitochondrial damage.

## Methods

### Reagents and tools table

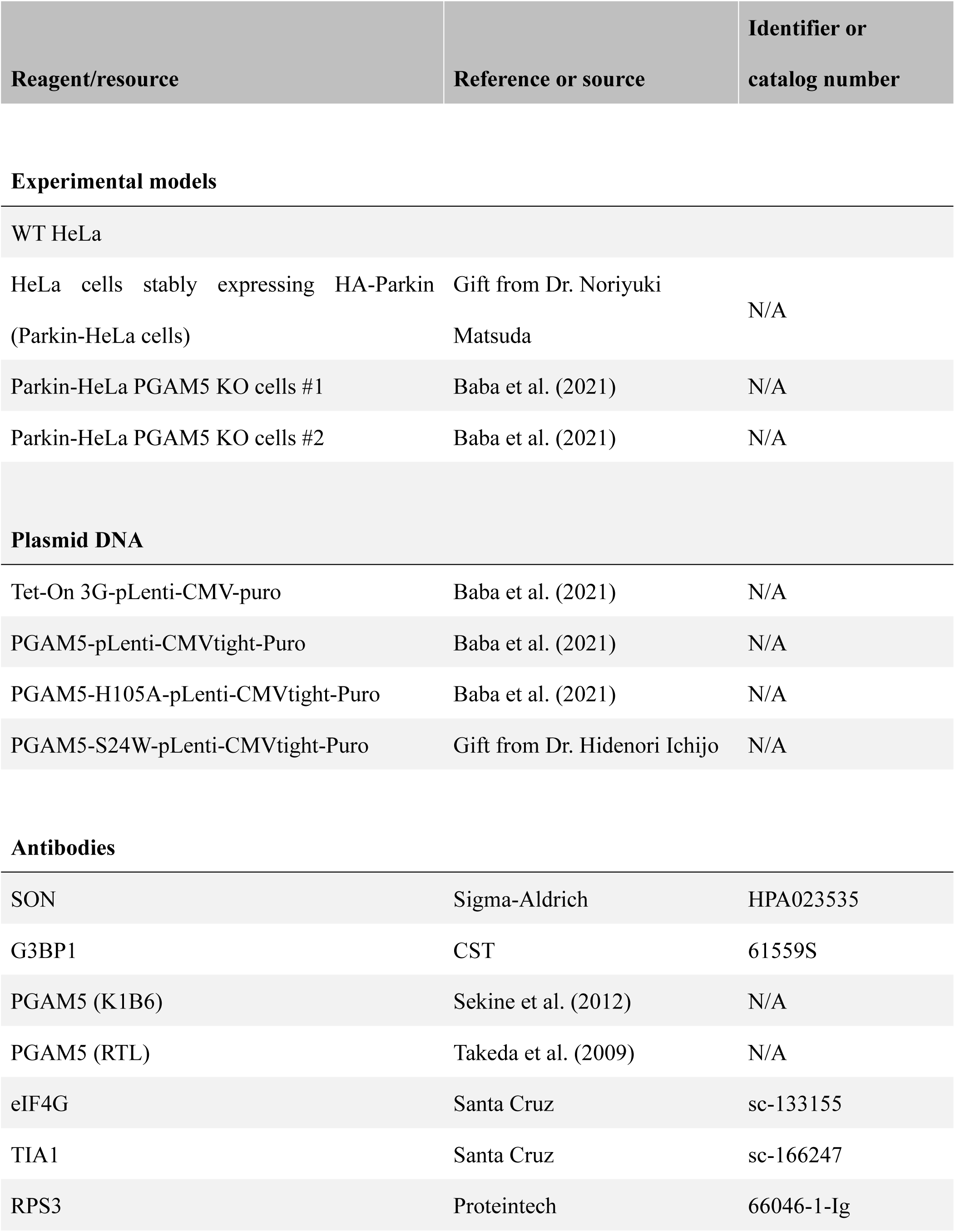
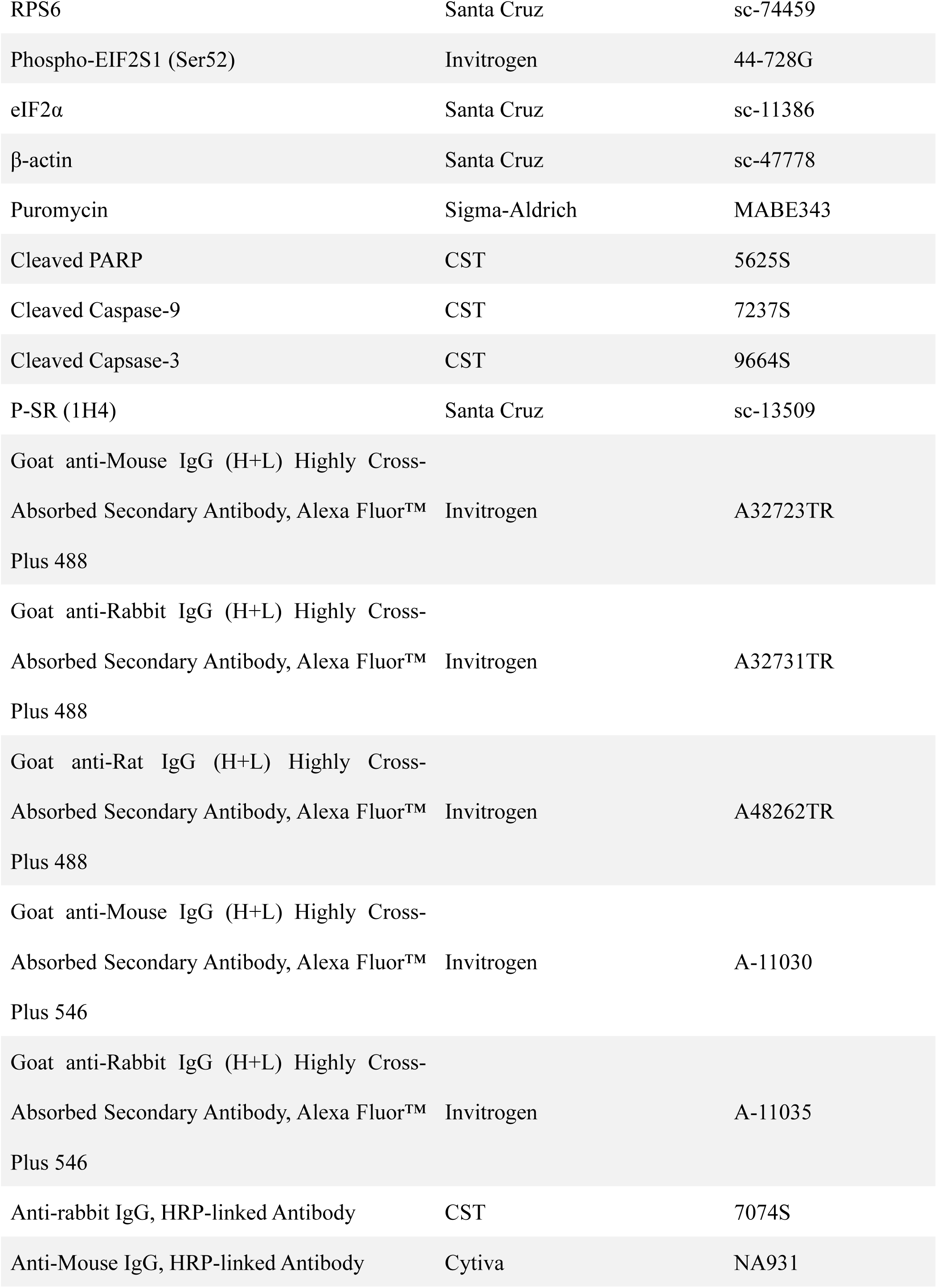
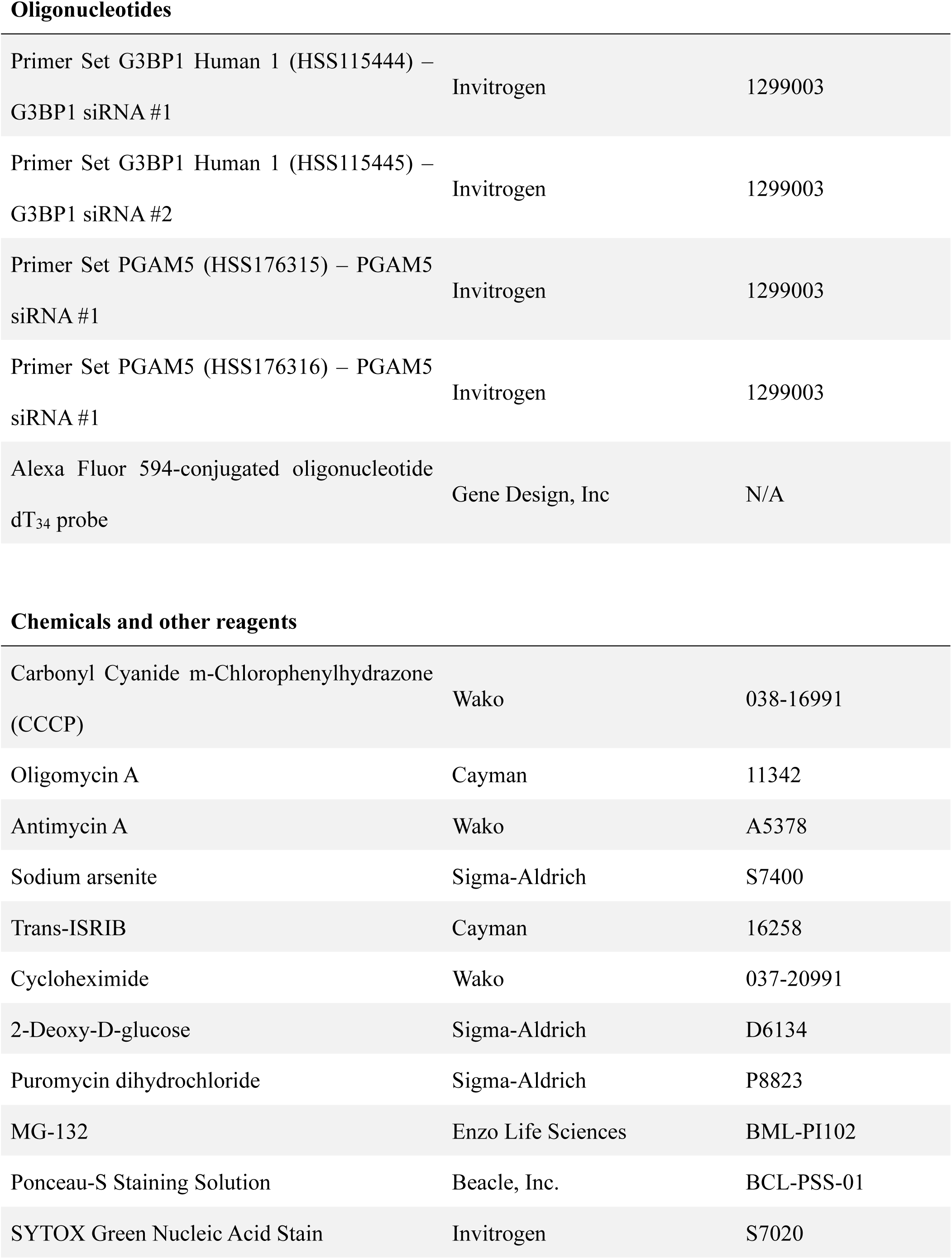
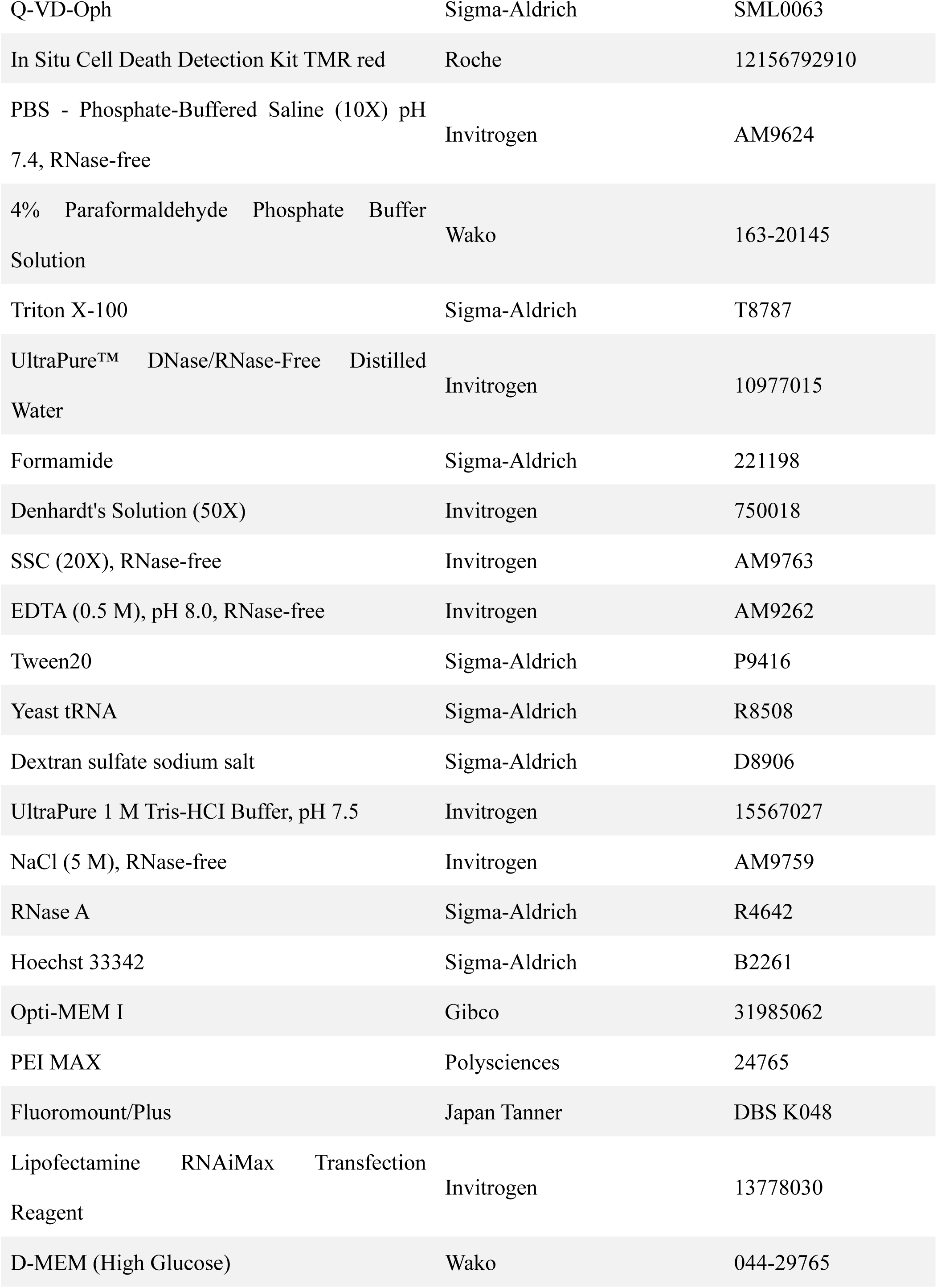
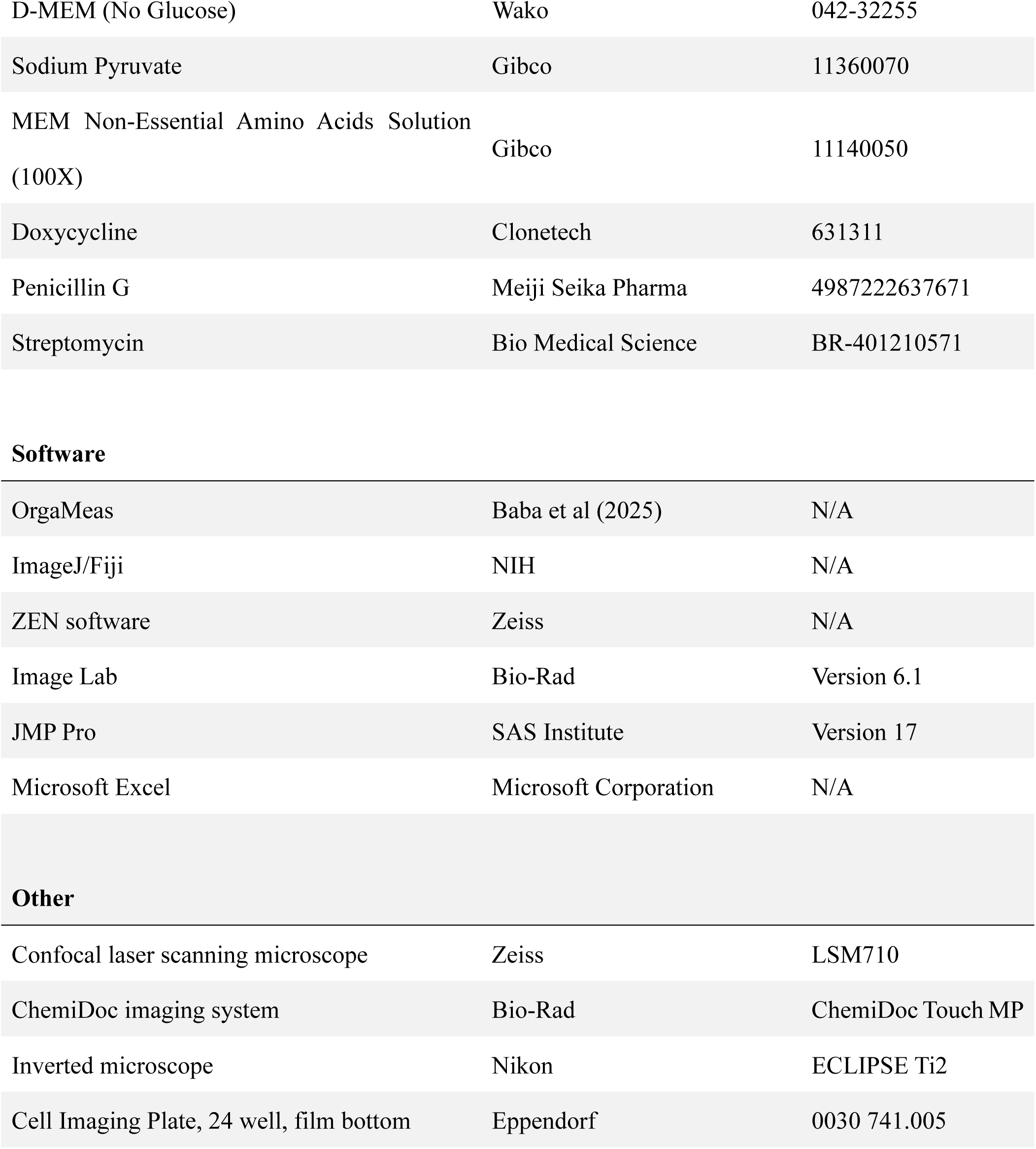

| Reagent/resource | Reference or source | Identifier or catalog number |
| --- | --- | --- |
| Experimental models |  |  |
| WT HeLa |  |  |
| HeLa cells stably expressing HA-Parkin (Parkin-HeLa cells) | Gift from Dr. Noriyuki Matsuda | N/A |
| Parkin-HeLa PGAM5 KO cells #1 | Baba et al. (2021) | N/A |
| Parkin-HeLa PGAM5 KO cells #2 | Baba et al. (2021) | N/A |
| Plasmid DNA |  |  |
| Tet-On 3G-pLenti-CMV-puro | Baba et al. (2021) | N/A |
| PGAM5-pLenti-CMVtight-Puro | Baba et al. (2021) | N/A |
| PGAM5-H105A-pLenti-CMVtight-Puro | Baba et al. (2021) | N/A |
| PGAM5-S24W-pLenti-CMVtight-Puro | Gift from Dr. Hidenori Ichijo | N/A |
| Antibodies |  |  |
| SON | Sigma-Aldrich | HPA023535 |
| G3BP1 | CST | 61559S |
| PGAM5 (K1B6) | Sekine et al. (2012) | N/A |
| PGAM5 (RTL) | Takeda et al. (2009) | N/A |
| eIF4G | Santa Cruz | sc-133155 |
| TIA1 | Santa Cruz | sc-166247 |
| RPS3 | Proteintech | 66046-1-Ig |
| RPS6 | Santa Cruz | sc-74459 |
| Phospho-EIF2S1 (Ser52) | Invitrogen | 44-728G |
| eIF2α | Santa Cruz | sc-11386 |
| β-actin | Santa Cruz | sc-47778 |
| Puromycin | Sigma-Aldrich | MABE343 |
| Cleaved PARP | CST | 5625S |
| Cleaved Caspase-9 | CST | 7237S |
| Cleaved Capsase-3 | CST | 9664S |
| P-SR (1H4) | Santa Cruz | sc-13509 |
| Goat anti-Mouse IgG (H+L) Highly Cross-Absorbed Secondary Antibody, Alexa Fluor™ Plus 488 | Invitrogen | A32723TR |
| Goat anti-Rabbit IgG (H+L) Highly Cross-Absorbed Secondary Antibody, Alexa Fluor™ Plus 488 | Invitrogen | A32731TR |
| Goat anti-Rat IgG (H+L) Highly Cross-Absorbed Secondary Antibody, Alexa Fluor™ Plus 488 | Invitrogen | A48262TR |
| Goat anti-Mouse IgG (H+L) Highly Cross-Absorbed Secondary Antibody, Alexa Fluor™ Plus 546 | Invitrogen | A-11030 |
| Goat anti-Rabbit IgG (H+L) Highly Cross-Absorbed Secondary Antibody, Alexa Fluor™ Plus 546 | Invitrogen | A-11035 |
| Anti-rabbit IgG, HRP-linked Antibody | CST | 7074S |
| Anti-Mouse IgG, HRP-linked Antibody | Cytiva | NA931 |
| **Oligonucleotides** |  |  |
| Primer Set G3BP1 Human 1 (HSS115444) –<br>G3BP1 siRNA #1 | Invitrogen | 1299003 |
| Primer Set G3BP1 Human 1 (HSS115445) –<br>G3BP1 siRNA #2 | Invitrogen | 1299003 |
| Primer Set PGAM5 (HSS176315) – PGAM5<br>siRNA #1 | Invitrogen | 1299003 |
| Primer Set PGAM5 (HSS176316) – PGAM5<br>siRNA #1 | Invitrogen | 1299003 |
| Alexa Fluor 594-conjugated oligonucleotide<br>dT<sub>34</sub> probe | Gene Design, Inc | N/A |
| **Chemicals and other reagents** |  |  |
| Carbonyl Cyanide m-Chlorophenylhydrazone<br>(CCCP) | Wako | 038-16991 |
| Oligomycin A | Cayman | 11342 |
| Antimycin A | Wako | A5378 |
| Sodium arsenite | Sigma-Aldrich | S7400 |
| Trans-ISRIB | Cayman | 16258 |
| Cycloheximide | Wako | 037-20991 |
| 2-Deoxy-D-glucose | Sigma-Aldrich | D6134 |
| Puromycin dihydrochloride | Sigma-Aldrich | P8823 |
| MG-132 | Enzo Life Sciences | BML-PI102 |
| Ponceau-S Staining Solution | Beacle, Inc. | BCL-PSS-01 |
| SYTOX Green Nucleic Acid Stain | Invitrogen | S7020 |
| Q-VD-Oph | Sigma-Aldrich | SML0063 |
| In Situ Cell Death Detection Kit TMR red | Roche | 12156792910 |
| PBS - Phosphate-Buffered Saline (10X) pH 7.4, RNase-free | Invitrogen | AM9624 |
| 4% Paraformaldehyde Phosphate Buffer Solution | Wako | 163-20145 |
| Triton X-100 | Sigma-Aldrich | T8787 |
| UltraPure™ DNase/RNase-Free Distilled Water | Invitrogen | 10977015 |
| Formamide | Sigma-Aldrich | 221198 |
| Denhardt's Solution (50X) | Invitrogen | 750018 |
| SSC (20X), RNase-free | Invitrogen | AM9763 |
| EDTA (0.5 M), pH 8.0, RNase-free | Invitrogen | AM9262 |
| Tween20 | Sigma-Aldrich | P9416 |
| Yeast tRNA | Sigma-Aldrich | R8508 |
| Dextran sulfate sodium salt | Sigma-Aldrich | D8906 |
| UltraPure 1 M Tris-HCl Buffer, pH 7.5 | Invitrogen | 15567027 |
| NaCl (5 M), RNase-free | Invitrogen | AM9759 |
| RNase A | Sigma-Aldrich | R4642 |
| Hoechst 33342 | Sigma-Aldrich | B2261 |
| Opti-MEM I | Gibco | 31985062 |
| PEI MAX | Polysciences | 24765 |
| Fluoromount/Plus | Japan Tanner | DBS K048 |
| Lipofectamine RNAiMax Transfection Reagent | Invitrogen | 13778030 |
| D-MEM (High Glucose) | Wako | 044-29765 |
| D-MEM (No Glucose) | Wako | 042-32255 |
| Sodium Pyruvate | Gibco | 11360070 |
| MEM Non-Essential Amino Acids Solution (100X) | Gibco | 11140050 |
| Doxycycline | Clontech | 631311 |
| Penicillin G | Meiji Seika Pharma | 4987222637671 |
| Streptomycin | Bio Medical Science | BR-401210571 |
| Software |  |  |
| OrgaMeas | Baba et al (2025) | N/A |
| ImageJ/Fiji | NIH | N/A |
| ZEN software | Zeiss | N/A |
| Image Lab | Bio-Rad | Version 6.1 |
| JMP Pro | SAS Institute | Version 17 |
| Microsoft Excel | Microsoft Corporation | N/A |
| Other |  |  |
| Confocal laser scanning microscope | Zeiss | LSM710 |
| ChemiDoc imaging system | Bio-Rad | ChemiDoc Touch MP |
| Inverted microscope | Nikon | ECLIPSE Ti2 |
| Cell Imaging Plate, 24 well, film bottom | Eppendorf | 0030 741.005 |

### Cell cultures

HeLa cells stably expressing HA-tagged Parkin (Parkin-HeLa cells) and PGAM5 KO Parkin-HeLa cells were established previously (Baba *et al*, 2021; Matsuda *et al*, 2010) and cultured in Dulbecco’s modified Eagle’s medium (DMEM) (high glucose; Wako, 044-29765) containing 8% fetal bovine serum (FBS), 100U/ml penicillin G (Meiji Seika Pharma, 4987222637671), 0.1 mg/ml streptomycin (Bio Medical Science, BR-401210571), 1 mM sodium pyruvate (Gibco, 11360070), 1× MEM non-essential amino acids (Gibco, 11140050), and 5 μg/ml puromycin (Sigma-Aldrich, P8823) under a 5% CO_2_ atmosphere at 37℃. HeLa cells were cultured in DMEM containing 8% FBS, 100 U/ml penicillin G, and 0.1 mg/ml streptomycin under a 5% CO2 atmosphere at 37°C.

### Expression plasmids

The expression plasmids, Tet-On 3G-pLenti-CMV-puro, PGAM5-pLenti-CMVtight-Puro, and PGAM5-H105A-pLenti-CMVtight-Puro, were constructed previously (Baba *et al*, 2021). PGAM5-S24W-pLenti-CMVtight-Puro was generated by the LR reaction based on Gateway recombinant cloning technology (ThermoFisher Scientific), in which PGAM5-S24W-pENTR was recombined with pLenti-CMVtight-Puro-DEST (a gift from Dr. Eric Campeau) (Campeau *et al*, 2009). Prior to that, PGAM5-S24W-pENTR was constructed by subcloning cDNA from PGAM5-S24W-pcDNA3 (gift from Dr. Hidenori Ichijo) into pENTER201.

### Plasmid DNA transfection

Expression plasmids were transfected into PGAM5 KO Parkin-HeLa cells using PEI MAX (Polysciences, 24765) according to the manufacturer’s instructions. PGAM5, PGAM5-H105A, and PGAM5-S24W were expressed in PGAM5 KO Parkin-HeLa cells by transfection with their respective expression plasmids and Tet-On 3G-pLenti-CMV-puro, followed by treatment with 25 ng/ml doxycycline (Clontech, 631311) for 24 h.

### siRNA transfection

Parkin-HeLa cells were transiently transfected with 20 nM final concentration of Stealth siRNA (ThermoFisher Scientific; HSS115444 and HSS115445 for G3BP1, and HSS176315 and HSS176316 for PGAM5) using Lipofectamine RNAiMax Transfection Reagent (Invitrogen, 13778030), according to the manufacturer’s instructions.

### Drug treatments and reagents

HeLa and Parkin-HeLa cells were treated with the following compounds/concentrations, unless otherwise indicated: CCCP 10 μM (Wako, 038-16991), oligomycin A 500 nM (Cayman, 11342), antimycin A 50 nM (Wako, A5378), sodium arsenite 0.25 mM (Sigma-Aldrich, S7400), Trans-ISRIB 0.5 μM (Cayman, 16258), cycloheximide 10 μg/ml (Wako, 037-20991), 2-deoxy-D-glucose 25 mM (Sigma-Aldrich, D6134), MG-132 5 μM (Enzo Life Sciences, BML-PI102), and Q-VD-Oph 10 μM (Sigma-Aldrich, SML0063). For glucose deprivation, cells were cultured in DMEM without glucose (No glucose; Wako, 042-32255) for 2 h.

### poly(A)^+^ RNA-FISH

Cells grown on glass coverslips in 24-well plates were washed with phosphate-buffered saline (PBS; Invitrogen, AM9624), fixed with 4% paraformaldehyde/PBS (Wako, 163-20145) for 15 min at room temperature, and then permeabilized with 0.5% Triton X-100 (Sigma-Aldrich, T8787) for 5 min at 4℃. The coverslips were washed with PBS and incubated in hybridization buffer (50% formamide (Sigma-Aldrich, 221198), 1× Denhardt’s solution (Invitrogen, 750018), 1× saline-sodium citrate (SSC; Invitrogen, AM9763), 10 mM ethylenediaminetetraacetic acid (EDTA; Invitrogen, AM9262), 0.01% Tween 20 (Sigma-Aldrich, P9416), and 0.1 mg/ml yeast tRNA (Sigma-Aldrich, R8508)) for 2 h at 55°C. An Alexa Fluor 594-conjugated oligonucleotide dT_34_ probe (Gene Design, Inc.) in hybridization buffer was added to the coverslips and incubated for 16 h at 55°C. The coverslips were washed with wash buffer (50% formamide, 2× SSC, and 0.01% Tween 20) for 15 min at 37°C and then incubated in 5 µg/ml RNase A (Sigma-Aldrich, R4642) in TNET buffer (50 mM Tris-HCl (Invitrogen, 15567027), pH 7.4, 140 mM NaCl (Invitrogen, AM9759), 5 mM EDTA, and 0.01% Triton X-100) for 30 min at 37°C. Subsequently, the coverslips were incubated in 2× SSC containing 0.01% Tween 20 for 15 min at 37°C and washed twice with 0.1× SSC containing 0.01% Tween 20 for 15 min at 37°C. After blocking with 1% bovine serum albumin (BSA) and 0.01% Tween 20 (Roche, 10735086001) in PBS for 1 h at room temperature, the coverslips were incubated in 1% BSA and 0.01% Tween 20 in PBS containing anti-G3BP1 (CST, 61559S) or anti-SON antibodies (Sigma-Aldrich, HPA023535) for 1 h at room temperature. After washing with PBS containing 0.2% Tween 20, a solution of 1% BSA and 0.01% Tween 20 in PBS containing Alexa Fluor 488-conjugated secondary antibody (Invitrogen, A32731TR, A48262TR) was added to the coverslips for 1 h at room temperature. After washing with PBS containing 0.2% Tween 20, the coverslips were incubated in PBS containing 20 μM Hoechst 33342 (Sigma-Aldrich, B2261) for 10 min at room temperature and mounted using Fluoromount/Plus reagent (Japan Tanner, DBS K048). Eight-bit images were acquired by confocal laser scanning microscopy (LSM710; Zeiss). To measure mRNA abundance in the intracellular compartments (Fig. S1A), the intensity of poly(A)^+^ RNA signals in each cell was quantified as the total signal intensity within the nuclear and cytoplasmic regions of the cells. The nuclear region was defined using a nuclear mask generated by OrgaSegNet, an organelle segmentation tool implemented in OrgaMeas (Baba *et al*, 2025). Cytoplasmic mRNA granules were identified by segmenting poly(A)^+^ RNA-positive foci using OrgaSegNet. Cells with two or more cytoplasmic foci were classified as cytoplasmic mRNA granule-positive cells. The proportion of cytoplasmic mRNA granule-positive cells and the size of the granules were quantified automatically at the single-cell level using OrgaMeas.

### Immunofluorescence microscopy

Cells grown on glass coverslips in 24-well plates were washed with PBS, fixed with 4% paraformaldehyde for 15 min, and then permeabilized with 0.25% Triton X-100 for 5 min at room temperature. After blocking with 2.5% BSA, the cells were stained with antibodies against the proteins of interest. Immune complexes were detected with Alexa Fluor 488- or 546-conjugated secondary antibodies (Invitrogen, A-11030; Invitrogen, A-11035) and the nuclei were counterstained with 20 μM Hoechst 33342 for 1 h at room temperature. The coverslips were mounted using Fluoromount/Plus reagent. Eight-bit images were acquired by confocal microscopy (LSM710; Zeiss). Both mitoRGs and SGs were identified by segmenting G3BP1- or TIA1-positive foci using OrgaSegNet. Cells containing two or more cytoplasmic foci were classified as mitoRG- or SG-positive cells. The proportion of mitoRG- or SG-positive cells was quantified automatically at the single-cell level using OrgaMeas. To count cleaved PARP-positive cells, the intensity of cleaved PARP signals in each cell was quantified as the mean signal intensity in the nuclear region (defined using a nuclear mask generated by OrgaSegNet). Cleaved PARP-positive cells were defined as cells with a mean signal intensity within the nuclear region ≥ 100, which accurately distinguished between cleaved PARP-positive and -negative cells. The proportion of cleaved PARP-positive cells was quantified automatically at the single-cell level using OrgaMeas.

### TUNEL assay

Cells grown on glass coverslips in 24-well plates were washed with PBS, fixed with 4% paraformaldehyde for 15 min, and then permeabilized with 0.25% Triton X-100 for 5 min at room temperature. Using an In Situ Cell Death Detection Kit, TMR red (Roche, 12156792910), TUNEL reaction mixtures were prepared on ice by mixing the enzyme solution (TdT) and label solution (TMR-dUTP) at a ratio of 1:9. A total volume of 20 μl of the reaction mixture was applied directly onto each coverslip and incubated for 1 h at 37°C. After incubation, the coverslips were washed three times with PBS and counterstained with 20 μM Hoechst 33342 for 20 min at room temperature. The coverslips were then washed twice with PBS, rinsed briefly with Milli-Q water, and mounted onto glass slides using Fluoromount/Plus reagent. Eight-bit images were acquired by confocal microscopy (LSM710; Zeiss). TUNEL-positive cells were counted by quantifying the intensity of the TUNEL signals in each cell as the mean signal intensity within the nuclear region (defined using a nuclear mask generated by OrgaSegNet). TUNEL-positive cells were defined as cells with a mean signal intensity ≥ 100, which accurately distinguished between TUNEL-positive and - negative cells. The proportion of TUNEL-positive cells was quantified automatically at the single-cell level using OrgaMeas.

### SYTOX green assay

Cells grown in a film-bottom 24-well plate (Eppendorf, 0030 741.005) were stimulated with 10 μM CCCP with or without 10 μM Q-VD-Oph for 12 h. The cells were further treated for 30 min with 0.5 μM SYTOX Green (Invitrogen, S7020), a high-affinity nucleic acid stain that easily penetrates cells with compromised plasma membranes, and the fluorescence was measured using a plate reader (Bio Tek, Winooski, VT, U.S.A.; Cytation3, Ex/Em = 488/523 nm).

### Immunoblot analysis

Cells were lysed in 1% Triton X-100 lysis buffer containing 25 mM Tris-HCl (pH 7.5), 150 mM NaCl, 5 mM EGTA, 1% Triton X-100, 5 μg/ml aprotinin, and 1 mM PMSF. After centrifugation at 21,500 × g for 15 min at 4℃, the supernatants were collected as cell lysates. The lysates were then fractionated by SDS-PAGE and electroblotted onto polyvinylidene difluoride membranes. The membranes were probed with primary and horseradish peroxidase (HRP)-conjugated secondary antibodies. Information on primary and secondary antibodies is provided in the Reagents and tools table. Protein bands were visualized using an enhanced chemiluminescence system and analyzed with ChemiDoc Touch MP (Bio-Rad). Band intensities were quantified using Image Lab Version 6.1 (Bio-Rad).

### Puromycin incorporation assay

Cells were treated with 20 μg/ml puromycin for 10 min immediately before cell lysis. Lysates were run via immunoblot and probed with puromycin antibody (see “Immunoblot analysis” section). Information on primary and secondary antibodies is provided in the Reagents and tools table.

### Statistical analysis

All quantitative data are presented as mean ± standard error (SE) from a minimum of three independent experiments. All statistics were calculated using JMP Pro Version 17 (SAS Institute). Statistical analyses were performed using Student’s t-test, Dunnett’s multiple-comparison test, or Tukey’s multiple-comparison test, as indicated in the figure legends. A P-value < 0.05 was considered statistically significant.

### Declaration of generative AI and AI-assisted technologies in the writing process

During the preparation of this manuscript, the authors used ChatGPT (ver 5.2) to assist with refining the wording and clarity of the text. Following the use of this tool, the authors carefully reviewed and edited the content as necessary and take full responsibility for the final content of the publication.

## Acknowledgements

We thank Dr. Noriyuki Matsuda for the gift of Parkin-HeLa cells, Dr. Hidenori Ichijo for PGAM5-S24W-pcDNA3, and Dr. Eric Campeau for pLenti-CMVtight-Puro-DEST. We also thank Dr. Mutsuhiro Takekawa and Dr. Hideki Nishitoh for their valuable advice and discussion, and Susan Furness, PhD, from Edanz (https://jp.edanz.com/ac) for editing a draft of this manuscript. This work was supported in part by the Platform Project for Supporting Drug Discovery and Life Science Research (Basis for Supporting Innovative Drug Discovery and Life Science Research (BINDS)) from the Japan Agency for Medical Research and Development (AMED) (JP25ama121032), JSPS KAKENHI (grant number JP23KJ1762 to T. Baba, JP21K06069 to S. Tanimura, and JP23K06117 to K. Takeda), the ANRI Fellowship (to T. Baba), and the Shionogi Infectious Disease Research Promotion Foundation (to S. Tanimura).

## Contributions

**Taiki Baba**: Conceptualization; Data curation; Formal analysis; Funding acquisition; Investigation; Methodology; Resources; Validation; Visualization; Writing–original draft; Writing–review and editing. **Akimi Inoue**: Investigation; Validation. **Yuya Nagahata**: Investigation; Validation. **Hibiki Tsutsumi**: Investigation; Validation. **Jun Takouda**: Investigation; Validation; Writing–review and editing. **Rena Onoguchi-Mizutani**: Methodology; Resources; Supervision. **Nobuyoshi Akimitsu**: Methodology; Resources; Supervision. **Susumu Tanimura**: Funding acquisition; Investigation; Validation; Writing–review and editing. **Kohsuke Takeda**: Conceptualization; Data curation; Funding acquisition; Project administration; Resources; Supervision; Writing–review and editing

---

**Figure S1.**
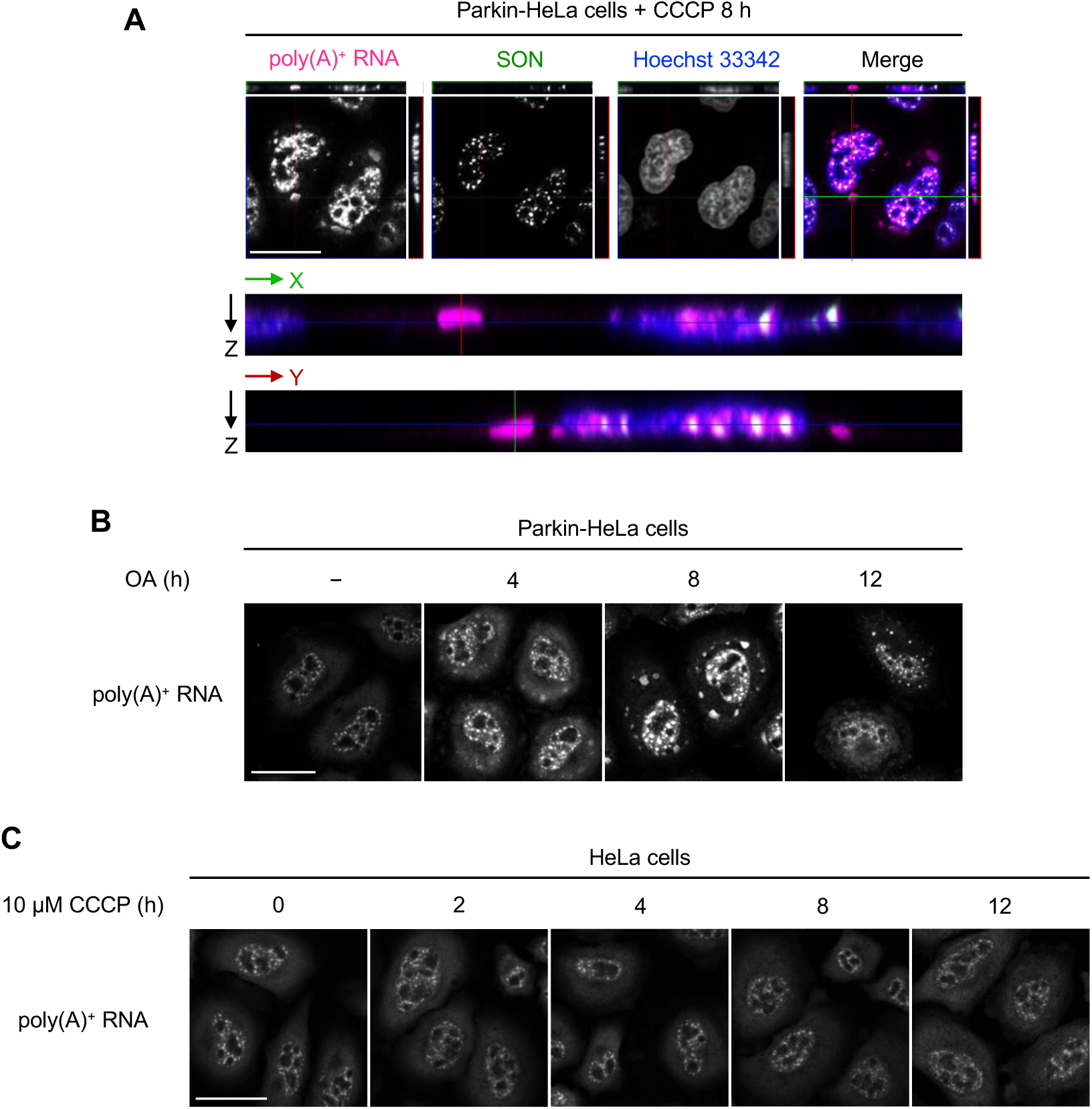
Mitochondrial stress induces the formation of cytoplasmic mRNA granules in a Parkin-dependent manner. (**A**) Z-stack of confocal images showing poly(A)^+^ RNA distribution. Parkin-HeLa cells treated with 10 μM CCCP for 8 h were subjected to poly(A)^+^ RNA-FISH, immunofluorescence analysis for SON as a nuclear speckle marker, and Hoechst 33342 staining (top panels). Z-stacks of merged confocal images from the XZ scans at the green line and YZ scans at the red line are shown in middle and bottom panels, respectively. (**B**) Representative results of poly(A)^+^ RNA-FISH in Parkin-HeLa cells treated with 500 nM oligomycin A and 50 nM antimycin A (OA) for 4, 8, and 12 h. Scale bar = 20 μm. (**C**) Representative results of poly(A)^+^ RNA-FISH in HeLa cells treated with 10 μM CCCP for 2, 4, 8, and 12 h. Scale bar = 20 μm.

**Figure S2.**
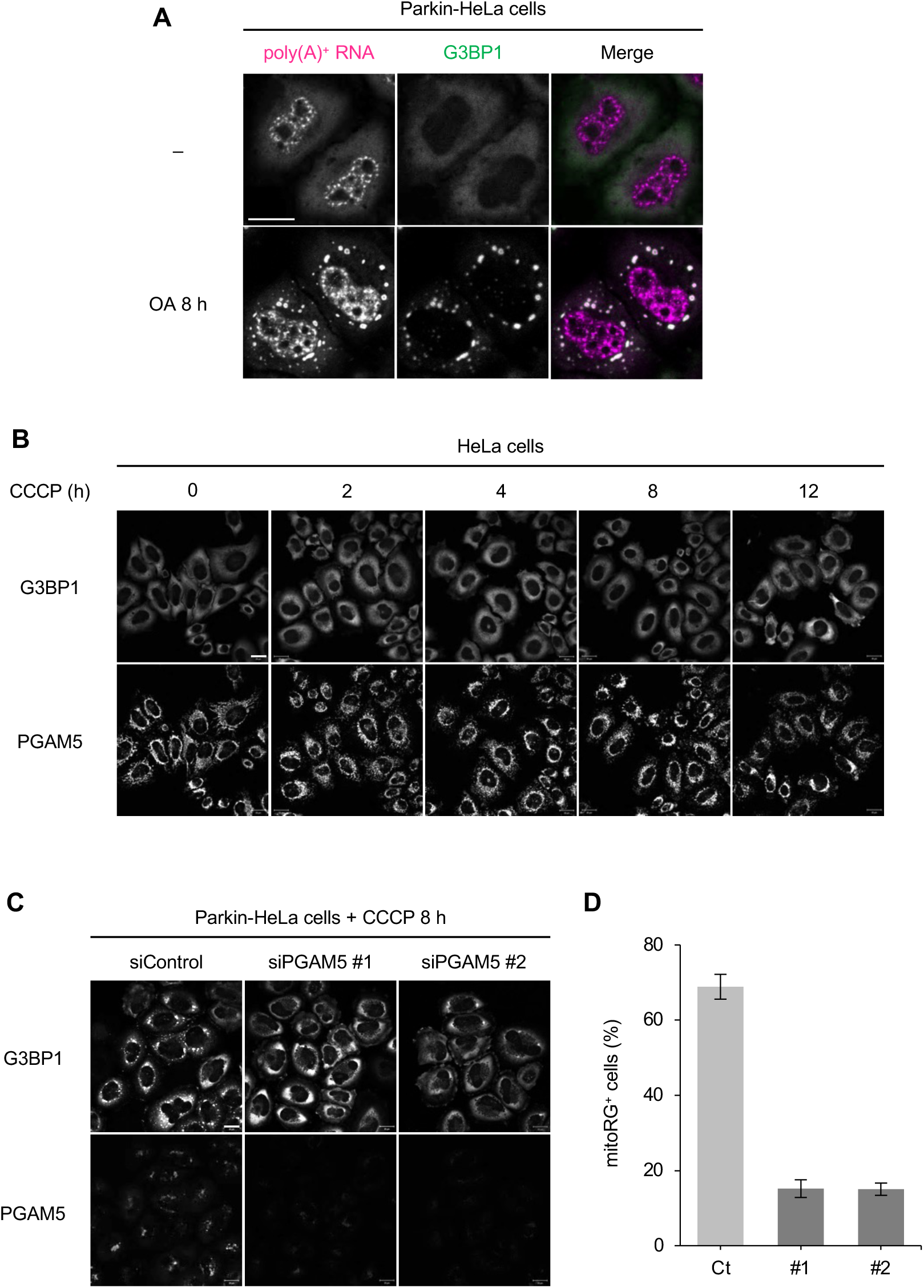
Mitochondrial stress induces mitoRG formation in a PGAM5-dependent manner during PINK1/Parkin mitophagy. (**A**) Representative results of poly(A)^+^ RNA-FISH and immunofluorescence analysis for G3BP1 in Parkin-HeLa cells treated with 500 nM oligomycin A and 50 nM antimycin A (OA) for 8 h. Scale bar = 20 μm. (**B**) Representative immunofluorescence results for G3BP1 and PGAM5 in HeLa cells treated with 10 μM CCCP for 2, 4, 8, and 12 h. Scale bar = 20 μm. (**C**) Parkin-HeLa cells transfected with siRNAs targeting PGAM5 (siPGAM5 #1 and #2) or control siRNA (siControl) were treated with 10 μM CCCP for 8 h and subjected to immunofluorescence analysis for G3BP1 and PGAM5. Scale bar = 20 μm. (**D**) Quantification of proportion of mitoRG^+^ cells in (C). Dunnett’s multiple-comparison test, ***P < 0.001. Error bars indicate mean ± SE. N = 3 independent experiments.

**Figure S3.**
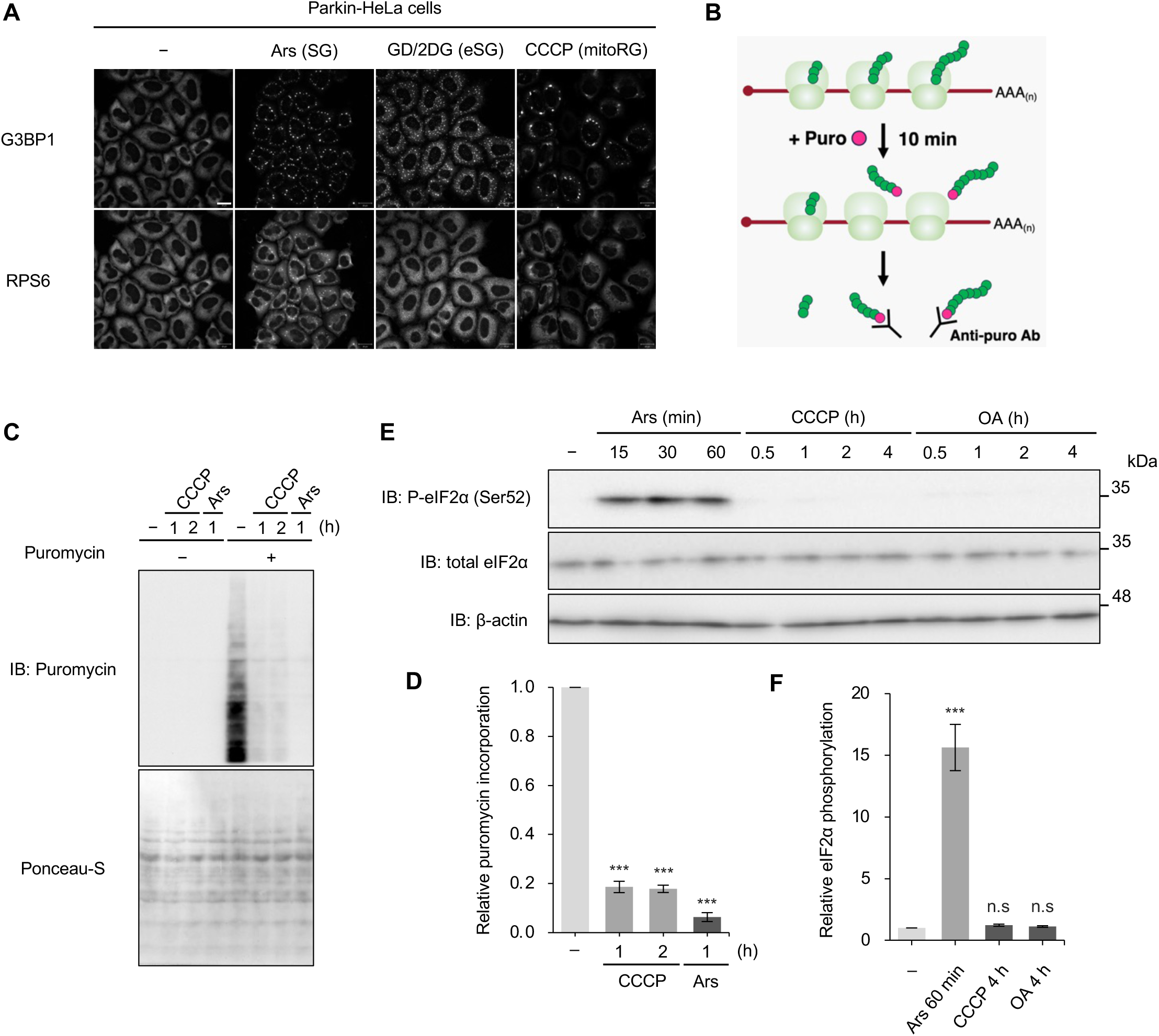
Global translation arrest occurs in the absence of eIF2α phosphorylation before mitoRG formation. (**A**) Representative immunofluorescence results for G3BP1 and RPS6 in Parkin-HeLa cells subjected to treatment with 0.25 mM sodium arsenite (Ars) for 1 h, glucose deprivation in combination with 25 mM 2-deoxy-D-glucose treatment for 2 h (GD/2DG), or treatment with 10 μM CCCP for 6 h, which induced SGs, energy deficiency-induced SGs (eSGs), or mitoRGs, respectively. Scale bar = 20 μm. (**B**) Outline of puromycin incorporation assay. (**C**) Parkin-HeLa cells treated with 10 μM CCCP or 0.25 mM Ars for the indicated periods were pulsed with 20 μg/ml puromycin for 10 min, followed by immunoblotting (IB) with puromycin antibody (upper panel). The total amount of loaded proteins was estimated by staining the membrane with Ponceau S (lower panel). (**D**) Relative puromycin intensity in (C) was quantified. Dunnett’s multiple-comparison test, ***P < 0.001. Error bars indicate mean ± SE. N = 3 independent experiments. (**E**) Parkin-HeLa cells treated with 0.25 mM Ars, 10 μM CCCP, or 500 nM oligomycin A and 50 nM antimycin A (OA) for the indicated periods were subjected to IB with the indicated antibodies. (**F**) Quantification of P-eIF2α levels in (E). Dunnett’s multiple-comparison test, ***P < 0.001. Error bars indicate mean ± SE. N = 3 independent experiments.

**Figure S4.**
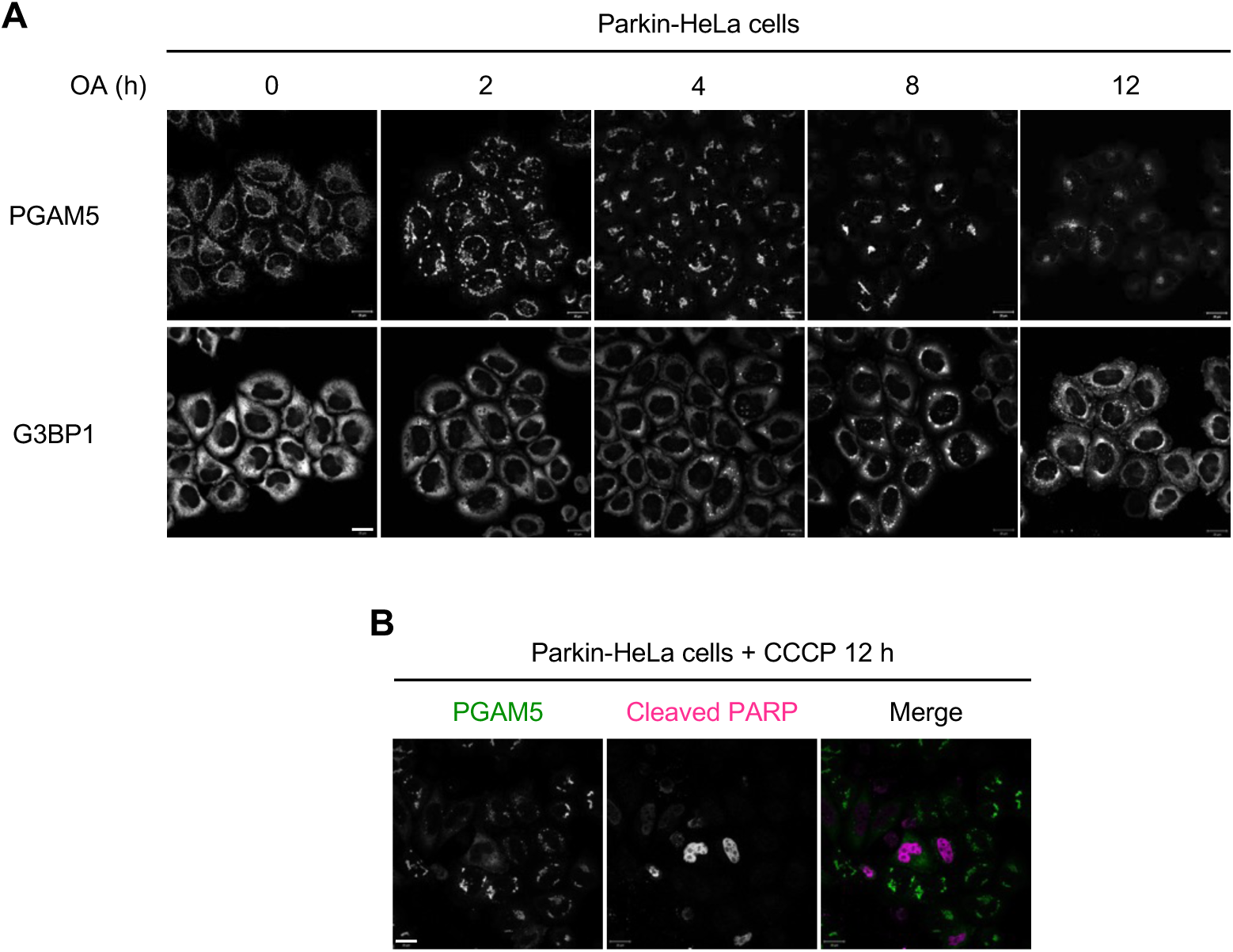
MitoRG disassembly and apoptosis induction correlate with the cytosolic diffusion of PGAM5 during PINK1/Parkin mitophagy. (**A**) Representative immunofluorescence results for PGAM5 and G3BP1 in Parkin-HeLa cells treated with 500 nM oligomycin A and 50 nM antimycin A (OA) for 2, 4, 8, and 12 h. Scale bar = 20 μm. (**B**) Representative immunofluorescence results for PGAM5 and cleaved PARP in Parkin-HeLa cells treated with 10 μM CCCP for 12 h. Scale bar = 20 μm.

**Figure S5.**
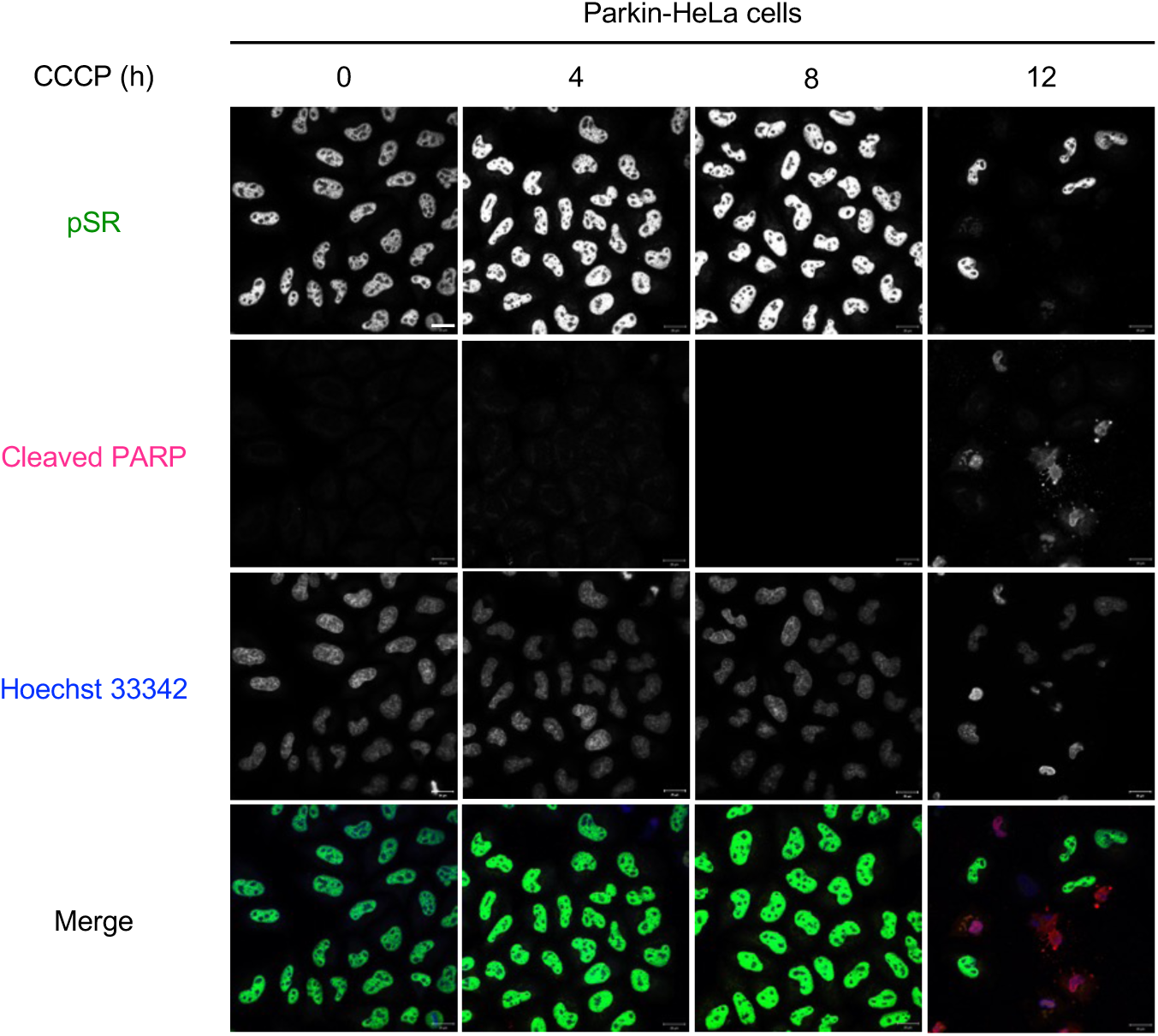
Decrease in SR protein phosphorylation levels correlates with apoptosis induction during PINK1/Parkin mitophagy. Representative immunofluorescence results for phosphorylated SR proteins, nuclear phospho-proteins involved in mRNA metabolism (Baba et al, 2021), and cleaved PARP in Parkin-HeLa cells treated with 10 μM CCCP for 4, 8, and 12 h, followed by counterstaining with Hoechst 33342. Scale bar = 20 μm.

